# A pan-tissue, pan-disease compendium of human orphan genes

**DOI:** 10.1101/2024.02.21.581488

**Authors:** Urminder Singh, Jeffrey A. Haltom, Joseph W. Guarnieri, Jing Li, Arun Seetharam, Afshin Beheshti, Bruce Aronow, Eve Syrkin Wurtele

## Abstract

Species-specific genes are ubiquitous in evolution, with functions ranging from prey paralysis to survival in subzero temperatures. Because they are typically expressed under limited conditions and lack canonical features, such genes may be vastly under-identified, even in humans. Here, we leverage terabytes of human RNA-Seq data to identify thousands of highly-expressed transcripts that do not correspond to any Gencode-annotated gene. Many may be novel ncRNAs although 80% of them contain ORFs that have the potential of encoding proteins unique to *Homo sapiens* (orphan genes). We validate our findings with independent strand-specific and single-cell RNA-seq datasets. Hundreds of these novel transcripts overlap with deleterious genomic variants; thousands show significant association with disease-specific patient survival. Most are dynamically regulated and accumulate selectively in particular tissues, cell-types, developmental stages, tumors, COVID-19, sex, and ancestries. As such, these transcripts hold potential as diagnostic biomarkers or therapeutic targets. To empower future discovery, we provide a compendium of these huge RNA-Seq expression data, and RiboSeq data, with associated metadata. Further, we supply the gene models for the novel genes as UCSC Genome Browser tracks.

## Introduction

Orphan genes – protein-coding genes unique to a given species- arise continually, providing organisms with a liminal reservoir of genetic elements for evolutionary innovation (1–6). Many orphan genes will lose utility and wane during the course of evolution; however, under persistent or intermittent environmental challenges, an orphan gene that improves survival will be retained, and its functionality optimized under selection (2, 4, 7, 8). A logical corollary is that genomes are composed of genes that arose at different evolutionary stages (1).

The evolutionary origins of orphan genes is varied. Orphan genes can form *de novo* from non-genic regions of the genome, introns, or by mitochondrial fostering; protein-coding orphan genes can also arise *de novo* from the emergence of novel reading frames within existing genes, antisense, gene fusions or other ncRNAs (3–12). Regardless of their ontogeny or whether they code proteins, newly emerged genes can be considered as a disruptive force in evolution, enabling remarkable traits to arise quickly in extant organisms (6).

Thousands of orphan genes from diverse species have been functionally characterized (2, 4, 6, 7, 13, 14); these represent only a tiny fraction of the estimated billions of the orphan genes in extant species (5).

Orphan genes encode paralyzing toxins of tens of thousands of species of jellyfish (2, 15) and parasitic wasps (16); “antifreeze” proteins, which have evolved independently in thousands of eukaryotic and prokaryotic species and enable survival in frigid temperatures (17); metabolic remodeling factors that mitigate pest and pathogen responses (13, 18); reshaping reproduction (19, 20); and functional or prognostic roles in context of human physiology and diseases (4, 14, 21, 22). The SARS-CoV-2 orphan gene, ORF10, enhances the clinical course of COVID-19 in part by attenuation of OXPHOS and innate immunity (23).

Empirical evaluation provides insight into the limitations of relying solely on standard *ab initio* approaches to identify orphan genes in a genome (5), likely because *ab initio* pipelines are biased towards canonical gene features and ubiquitous expression, neither of which are characteristics of orphan genes (5, 7, 24). Incorporating direct evidence from high-throughput data can identify novel genes that are undetected or discarded by other approaches (5, 7, 11, 25–27). To capture those genes expressed under limited conditions, it is critical to use RNA-Seq data from diverse tissues, populations, diseases and treatments (5).

Even when described in the literature, novel transcripts are rarely integrated into community databases (4, 5, 21, 28) and thus are difficult to access for further study. Furthermore, even if annotated in a community database, transcripts/genes not designated as “protein-coding” are often intentionally excluded from consideration in ongoing research.

Here, we identify novel human transcripts on a massive scale, validate those that are highly-expressed in data from independent studies, and characterize them by their inferred phylostratal and genomic context.

## Results

### Identification of novel evidence-based (EB) transcripts

To optimize retrieval of novel human transcripts, while reducing study-specific biases (29, 30), we developed a reproducible workflow for uniform and harmonized read alignment, transcript assembly, and quantification of aggregated RNA-Seq data (Figure 1 A). The workflow is carefully documented to enable its efficient reuse by researchers on other datasets. Using this workflow, we re-processed raw reads from 26,985 RNA-Seq paired-end samples from Gene-Tissue Expression portal (GTEx) (31) and The Cancer Genome Atlas (TCGA; https://www.cancer.gov/tcga).

**Fig. 1.**
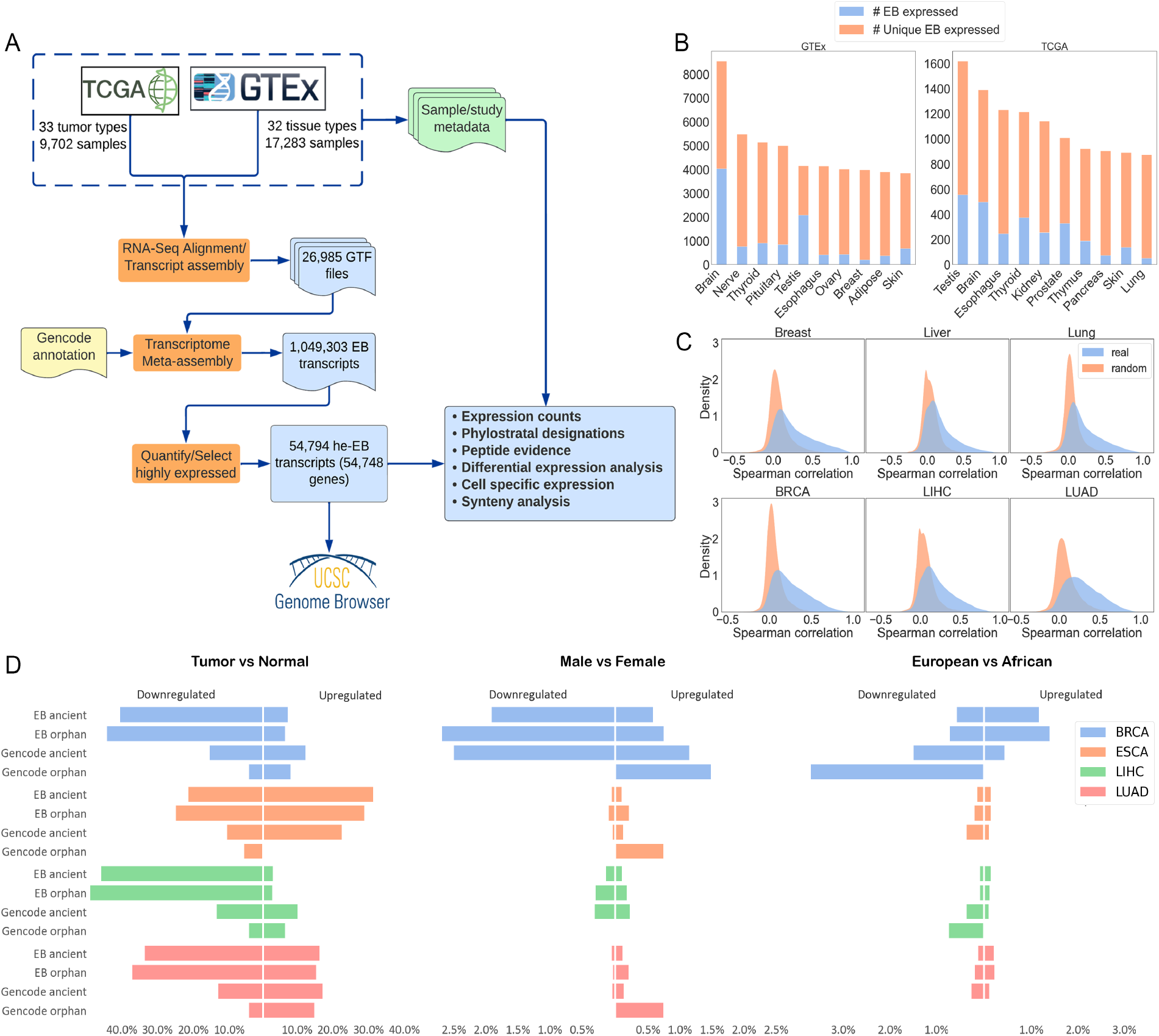
An overview of the orphan gene annotation workflow and expression patterns of the EB genes. **(A)** 26,985 RNA-Seq datasets were downloaded from TCGA and GTEx and each was processed through our alignment and transcript assembly pipeline. The 26,985 annotations were merged into a single consolidated annotation with our meta-assembly pipeline (see Fig. 13 for details). 54,794 EB transcripts with median expression comparable to protein-coding transcripts were retained for further analysis. **(B)** Number of highly expressed EB transcripts according to tissue for the GTEx and TCGA cohorts. 151 TCGA-LAML samples expressed 29,631 EB transcripts (Supplementary File 1). **(C)** Distributions of Spearman’s correlation values between each intronic EB transcript and its corresponding annotated transcript versus between random pairs of intronic and annotated transcripts. **(D)** Percentage of annotated and EB, orphan and ancient genes that are differentially expressed in four tumor tissues, between disease, sex, and self-reported race.

We merged the resulting 1,049,303 novel EB transcripts with the 335,239 transcripts of all types that are annotated in Gencode. The median expression of the EB transcripts was similar to that of the Gencode-designated lncRNAs, but less than that of protein-coding transcripts. We evaluated the expression levels of all novel and annotated transcripts from each tissue and tumor in a sex-, age- and race-specific manner, and merged these together for each tissue or tumor. This approach enabled us to retain those genes that are highly expressed only in individuals of a particular sex or ancestry. We minimized false positives (“noise”) by choosing a very stringent expression criteria for inclusion of a transcribed sequence to be considered as potential novel genes; we retained only novel EB transcripts with a median expression in a given tissue or tumor equal to or greater than that of the median of all protein-coding transcripts. Thus, many of the novel EB transcripts we detected, but did not include in our more detailed analysis, were expressed to the level of many annotated transcription factors. Only 54,794 of the over one million expressed but unannotated transcripts exceeded this level of accumulation; we term these high-expression EB transcripts (heEBs). Of these 54,794, 52,032 contained ORFs of over 99 nt.

### Thousands of novel heEB transcripts are expressed dynamically in diseased and undiseased tissues

Somewhat counter-intuitively, because cancer ontogeny is associated with mutation (32, 33), many times more heEB genes are expressed in non-diseased tissues than in the corresponding tumors (Figure 1 B).

Only 59 of the heEB genes are highly expressed in all 39 undiseased tissues analyzed from GTeX (Supplementary File 1); 59 are expressed in all of the tumor types; 35,212 are expressed in at least one tumor type, while 8,607 EB transcripts are expressed in both an undiseased tissue *and* a tumor in the same tissue-type.

We assessed the genomic locations of heEB genes with respect to the reference genome. The majority of heEB transcripts (84%) are located within introns of annotated genes (“intronic”), while 26% of the transcripts are located between annotated genes (“intergenic”).

Some EB transcripts located in introns might represent processing errors, e.g., premature termination, intron retention, or intronic reads, though careful studies indicate this is not the case for many novel intronic transcripts (34, 35). Reasoning that any intronic EB transcripts that were highly coexpressed with its subtending annotated transcript might be processing artifacts, we computed Spearman correlations for each of these pairs. However, the vast majority of EB transcripts are independently expressed (Figure 1 C; Supplementary File 1).

To identify orphan and primate-specific genes, the phylostrata of all heEB transcripts containing ORFs over 99 nt, and all Gencode protein-coding transcripts were evaluated by *phylostratr* (36). We extended the search space using Liftoff (37) to cover protein-coding regions that might be present but not yet annotated in target species, mapping human heEB genes to the whole genomes of nine closely related species, and based final phylostratal assignments on *phylostratr* (36) predictions merged with Liftoff designations (Figure 2). 13,921 of the heEB transcripts predicted by *phylostratr* as orphans had corresponding ORFs in apes (Hominoidea-specific), while 29,813 of the heEB transcripts were inferred to be orphan genes, found only in *Homo sapiens*.

**Fig. 2.**
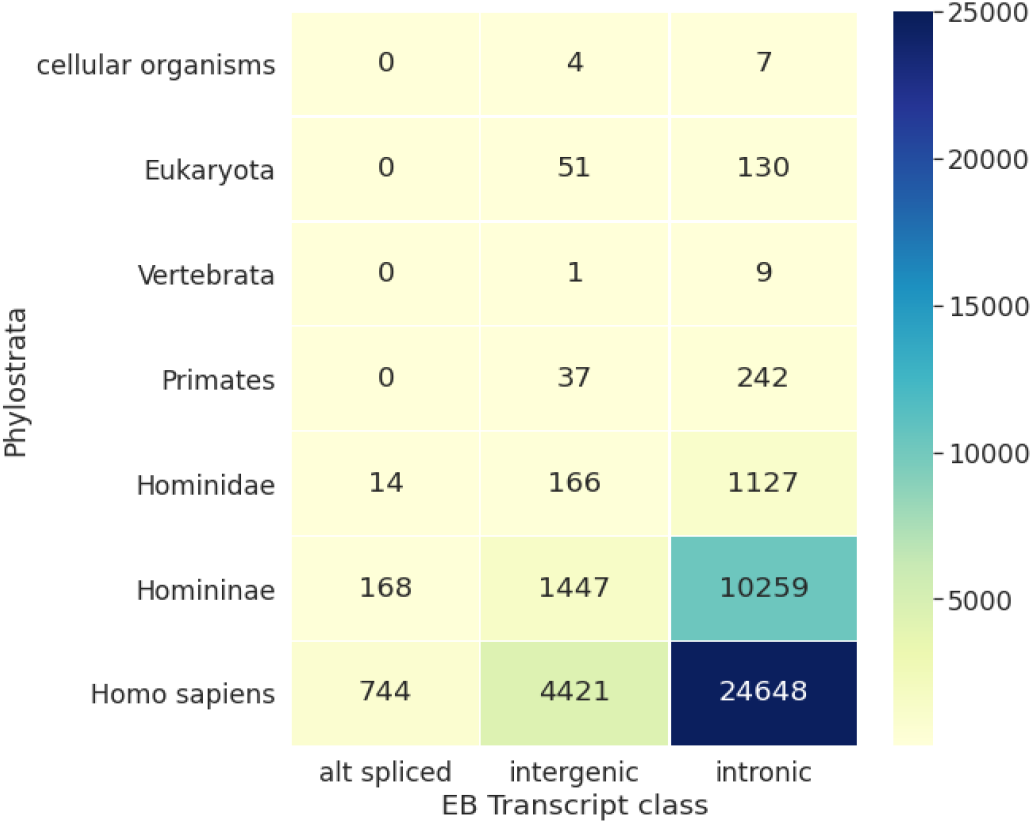
Phylostratal assignments and genomic contexts of novel heEB and anno-tated proteins-coding transcripts. Gene assignments combined predictions from phylostratr and Liftoff.

We hypothesized that as-yet unidentified heEB transcripts would be induced in COVID-19. These transcripts might have relevance as biomarkers of the viral disease, or as pharmaceutical targets. To investigate this, we utilized COVID-19-related RNA-Seq data from SRA. We selected 2,264 RNA-Seq samples from 32 COVID-19 studies (including the non-SARS-CoV-2 infected controls from these studies) and their corresponding sample metadata https://github.com/jahaltom/Orphan-Gene-Supp. Ancestry was assigned to a portion of the data using RIAD (38). The studies included samples obtained from nasopharyngeal tissue, blood cells and other in-vivo tissue, autopsied organs, as well as organoids. These data were mapped to the (human transcriptome (GencodeV36), SARS-COV-2 transcriptome (ASM985889v3), and the 1,049,303 human EB transcripts. We evaluated the expression levels of all transcripts from each tissue in a study/COVID-status/ancestry-specific manner, (sample size of >=5), using the same protocol as for the TCGA and GTeX data sets, with median TPM requirement of >=Q75 of the medians of all Gencode-identified protein coding transcripts.

This approach revealed 8,865 EB transcripts that are highly expressed in tissues infected by SARS-CoV-2, but not in corresponding control tissues; of these, 5,834 are orphans (Figure 3). As expected from observations on the TCGA/GTEx data, diverse SRA samples show distinct EB expression patterns.

**Fig. 3.**
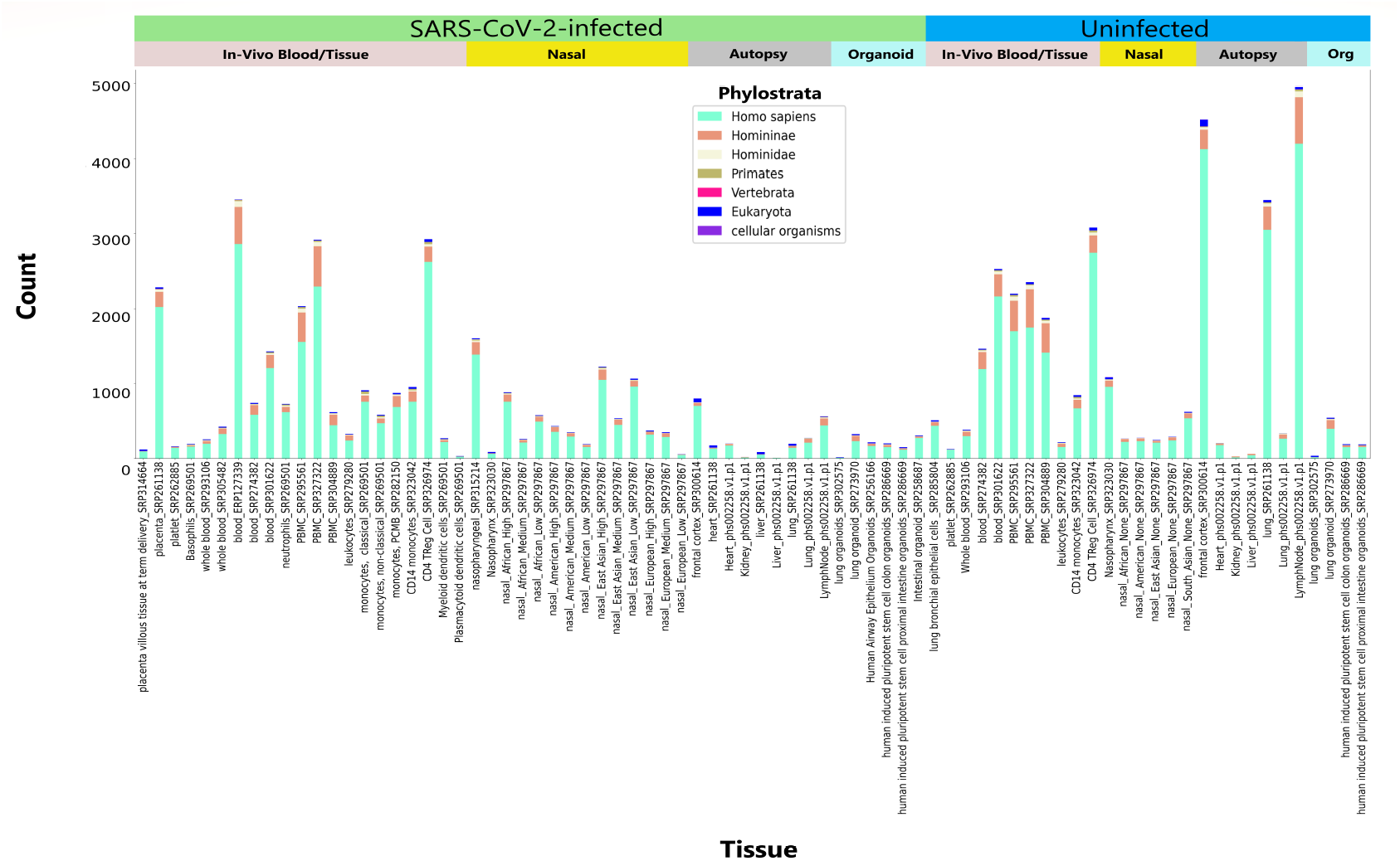
Expression of heEB genes in diverse SARS-CoV-2-infected and uninfected samples. Twenty seven studies https://github.com/jahaltom/Orphan-Gene-Supp with samples from nasal swabs, autopsy tissues, and in-vivo blood/tissues from individuals with COVID-19, and *in vitro* SARS-CoV-2-infected cultured organoids, as well as associated controls. Tissues were split into study/COVID-status/ancestry specific manner, each as a condition (x-axis). Count represents the number of highly expressed EB genes in a given condition. To be considered as highly expressed, an EB gene must have had a median TPM greater or equal to the Q75 of the medians of all known protein coding genes. Each condition has at least a sample size of five. The nasal swab conditions (High, Median, Low, None) depict viral load and are from study SRP297867.

### Novel genes exhibit differential expression across disease, sex, and ancestry

We identified differentially-expressed (DE) genes between tumor samples and corresponding undiseased samples, controlling for age, sex, and race.

Thousands of heEB genes are DE between diseased and undiseased samples. Notably, the upregulated and downregulated genes varies with tissue and disease type, self reported race, and sex. Figure 1 D exemplifies this finding, showing the percentages of heEB and annotated genes DE for four tumor types compared to the corresponding undiseased tissues.

19,201 heEB orphan genes are downregulated in THCA, whereas only 6,953 such genes are downregulated in KIRK. 8,576 heEB orphan genes are upregulated in ESCA, but only 849 heEB orphan genes are upregulated in THCA. 305 heEB orphans are downregulated and 11 are upregulated in common across every comparison between a tumor and its undiseased counterpart (Supplementary File 1).

Thousands of the EB orphan genes are DE by sex and ancestry (Figure 1 D). In PRAD, 670 heEB orphan genes are upregulated in African-Americans as compared to European-Americans; in BRCA, 416 genes are upregulated in European-Americans compared to African-Americans. Due to insufficient metadata, it cannot be determined whether such differences are a consequence of genetic or socio-environmental factors (39); regardless, they are an important consideration.

We validated our results using an independent study of strand-specific RNA-seq data comprising paired samples from 25 COAD tumors and adjacent undiseased colon tissue from African-American individuals (40). Sixty-six percent of the heEB genes that are upregulated in the COAD samples from TCGA are also upregulated in the independent COAD study, and 73% of the heEB genes downregulated in the COAD TCGA samples are also downregulated in COAD the independent COAD study (Supplementary File 1).

To validate the expression of heEB transcripts identified from GTEx and TCGA, we reprocessed raw data from two independent strand-specific RNA-Seq studies, which resolves ambiguous reads in overlapping genes transcribed from opposite strands, provides more accurate expression counts and ascertains intronic monoexonic transcripts from the antisense strand (35, 41). In eight samples from human liver, heart, brain, and testis, 1,690 EB transcripts are expressed. In 50 RNA-Seq samples from colon tumors and adjacent undiseased tissues, 1,803 EB transcripts are expressed (Supplementary File 1). The majority of the heEB transcripts from both these datasets potentially code for a human-specific protein (Figure 4 C and D).

**Fig. 4.**
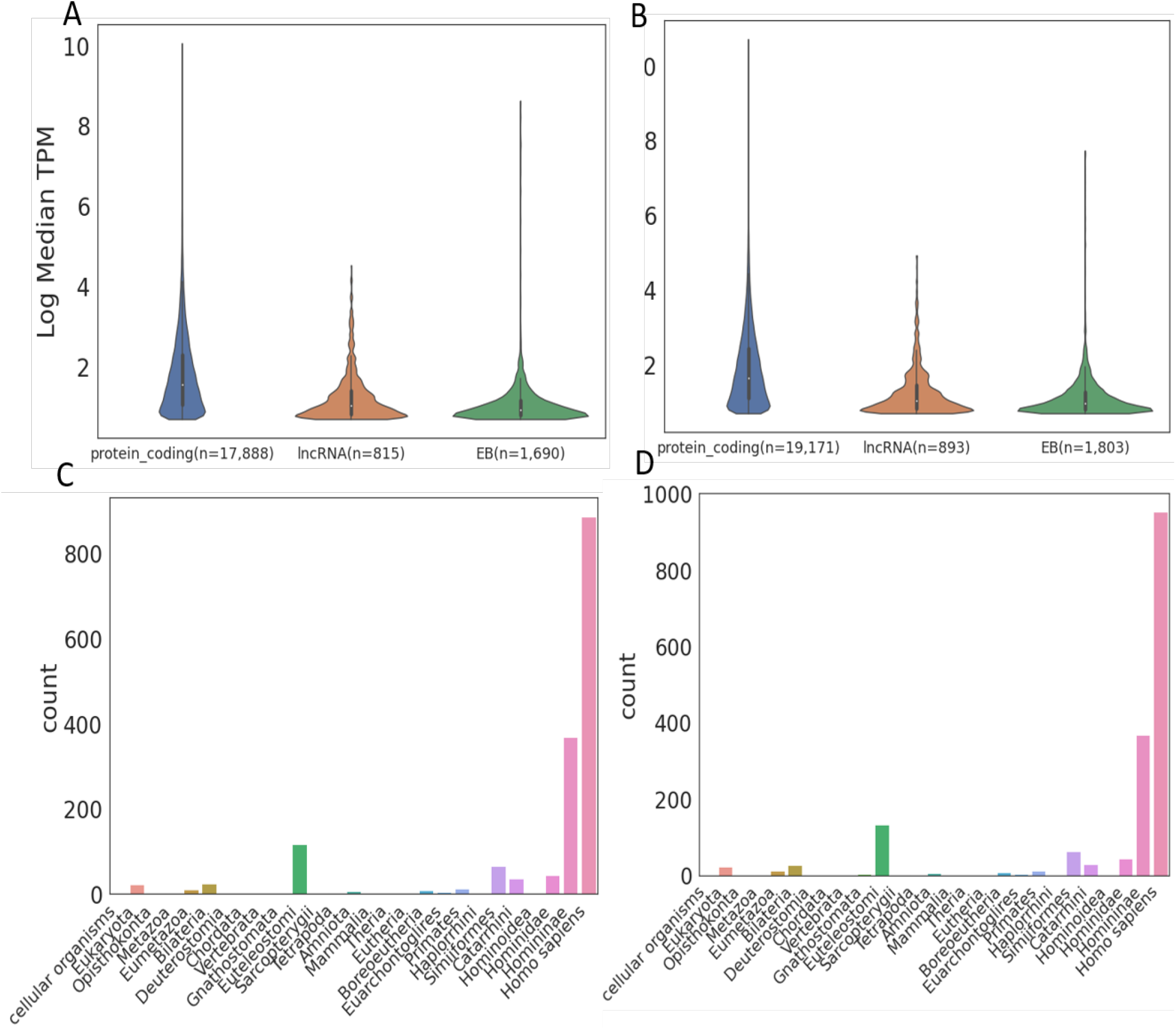
Expression of annotated protein-coding and lncRNA, and EB transcripts in independent, strand-specific RNA-Seq datasets. **A**. Eight RNA-Seq samples from liver, heart, brain, and testis of two individuals; 1,690 heEB transcripts were detected. **B**. Fifty RNA-Seq samples from colon cancer and adjacent undiseased tissues; 1,803 heEB transcripts were detected. Eight hundred and ninety heEB transcripts are expressed in both datasets. **C**. Frequency of phylostratal assignments for the 1,690 heEB transcripts expressed in samples from **A. D**. Frequency of phylostratal assignments for 1,803 heEB transcripts expressed in samples from **B**.

11,510 heEB transcripts have an ORF that is also present in the chimpanzee genome. We used eight chimpanzee stranded RNA-Seq samples from liver, heart, brain, and testis (35) to assess the expression of these heEB transcripts in this close human relative. 564 human heEB genes were expressed in the chimpanzee samples (median TPM *>*= 1) (Figure 5; Supplementary File 1).

**Fig. 5.**
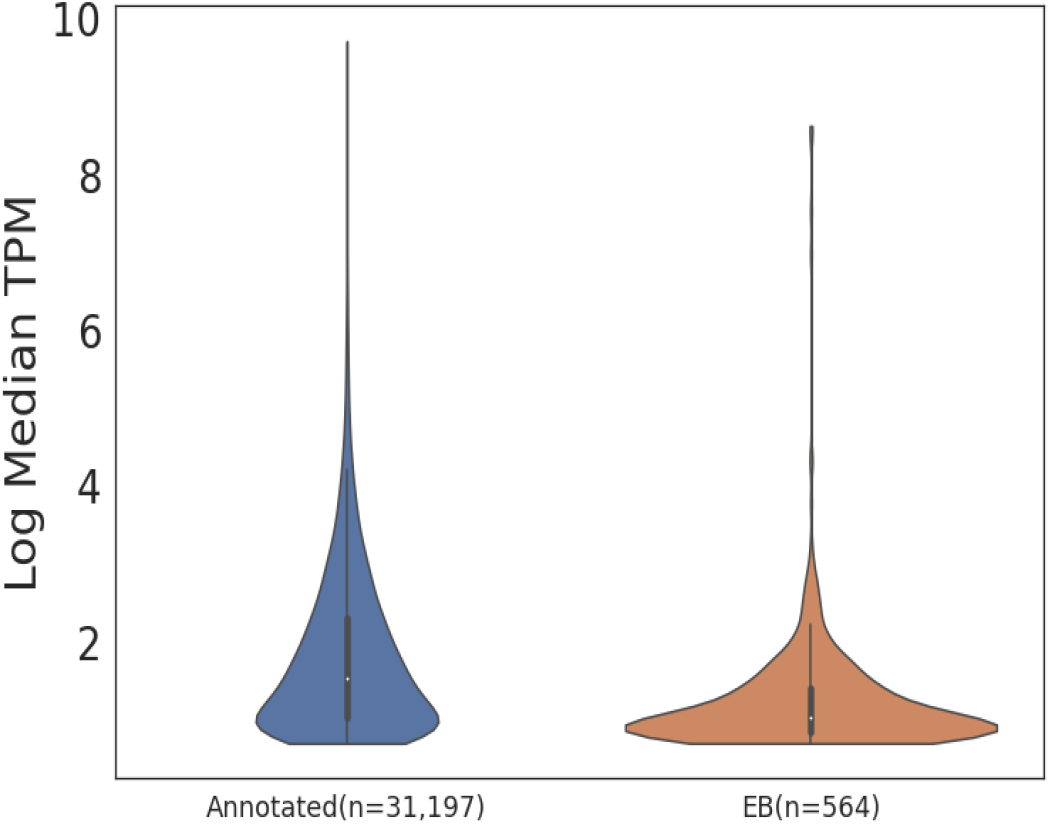
Expression distribution of Gencode-annotated and heEB orphan gene transcripts in chimpanzee strand-specific RNA-Seq data. The data consists of 8 RNA-Seq samples from chimpanzee liver, heart, brain, and testis. Five hundred and sixty-four EB transcripts (only 5% of the 11,510 unannotated transcripts that we quantified) were detected as expressed in chimpanzee.

### Cell- and developmental-specific expression of novel genes

To examine whether/which heEB genes are regulated in a cell-specific manner, we used independent scRNA-Seq datasets from liver, breast, testis, and lung, and identified cell-clusters corresponding to different cell-types (Supplementary Fig. 20). EB genes are among the top markers of cell-type clusters in breast, liver, and testis. Fig. 6 shows selected EB genes that mark particular clusters and have evidence of translation in Ribo-Seq datasets. Because of the cell- and condition-specific expression of many heEB genes, we anticipate that some may provide prognostic markers of the various unique cell-types.

**Fig. 6.**
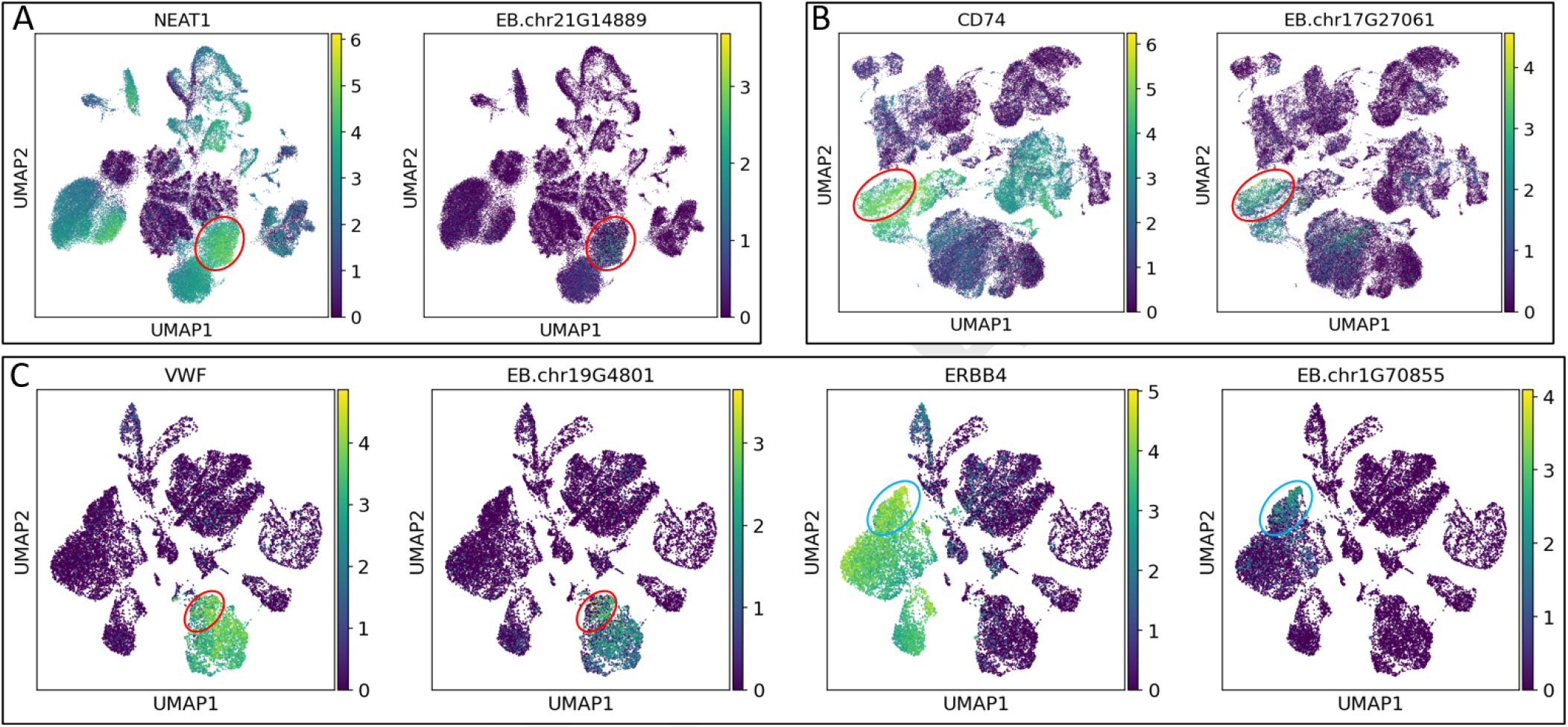
Cell-specific expression of novel genes. (A-C) Reprocessed raw data from single-cell RNA-Seq. UMAP visualizations (Supplementary Fig. 20) to identify expression patterns for heEB genes with Ribo-Seq evidence of translation (right) along with an annotated marker gene of the cluster (left). **A**. Adult liver. EB.chr21.G14889 is a marker gene for Cluster 3 (circled in red). **B**. Adult breast. We identified 45 clusters. The novel gene, EB.chr17.G27061 is a marker gene for Cluster 11 (circled in red). **C**. Adult testis. We identified 31 clusters. EB.chr19G4801 and EB.chr1G70855 show cluster-specific expression in two different clusters. EB.chr19G4801 co-localizes with the signature genes of Cluster 8 (endothelial cells) (circled in red). EB.chr1G70855 is highly expressed in Cluster 2, a subset of sertoli cells (circled in blue), a subset of sertoli cells.

A recent study (42) has provided unprecedented views of cell-types and cell sub-types of mammalian lungs in multiple species and across stages of development. These data thus offer the opportunity to identify novel transcripts and the predicted proteins associated with each. We re-aligned single nuclei RNA-Seq data derived from fetal, child, and adult lung samples. Leveraging cell annotations learned from prior analyses of human lung cells, we derived a database of cell signatures that could be used to identify cell-type and developmental stage-based gene signatures. All novel heEB genes and Gencode-annotated genes were co-segregated and evaluated along with the known cell-, class- and subclass-specific gene signatures. Those novel heEB genes that we detected as cell-specific are all inferred to be orphans (Fig. 7). The resultant, expanded, cell-specific or cell-preferential signatures, which are derived from known genes and heEB or Gencode genes of unclear function, are visualized in Supplementary Figure 1 and provided in the Human Cell Atlas project (https://software.broadinstitute.org/morpheus) for interactive exploration. This approach allows for the functional inferences of uncharacterized genes by leveraging knowledge associated with genes of known function.

**Fig. 7.**
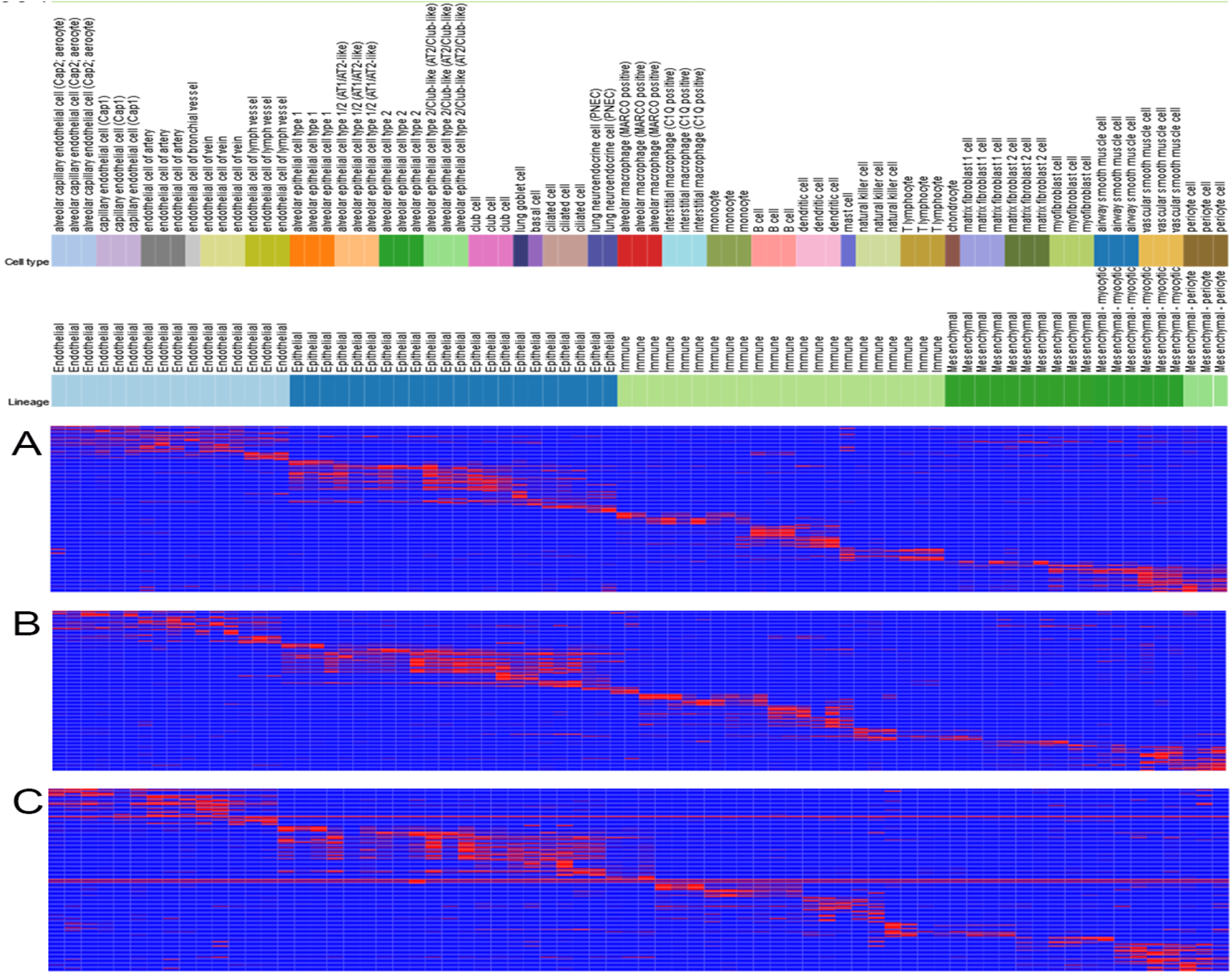
Cell-type-specific and developmental-stage-specific expression of novel orphan genes in human lungs. Raw data from single nuclei RNA-Seq of lung samples from fetal, child and adult individuals was processed to quantify expression of Gencode-identified and heEB genes. Visualized in Morpheus. Cell-type expression patterns of orphan heEB genes in **(A)** fetal **(B)** children **(C)** adult lung samples.

**Fig. 8.**
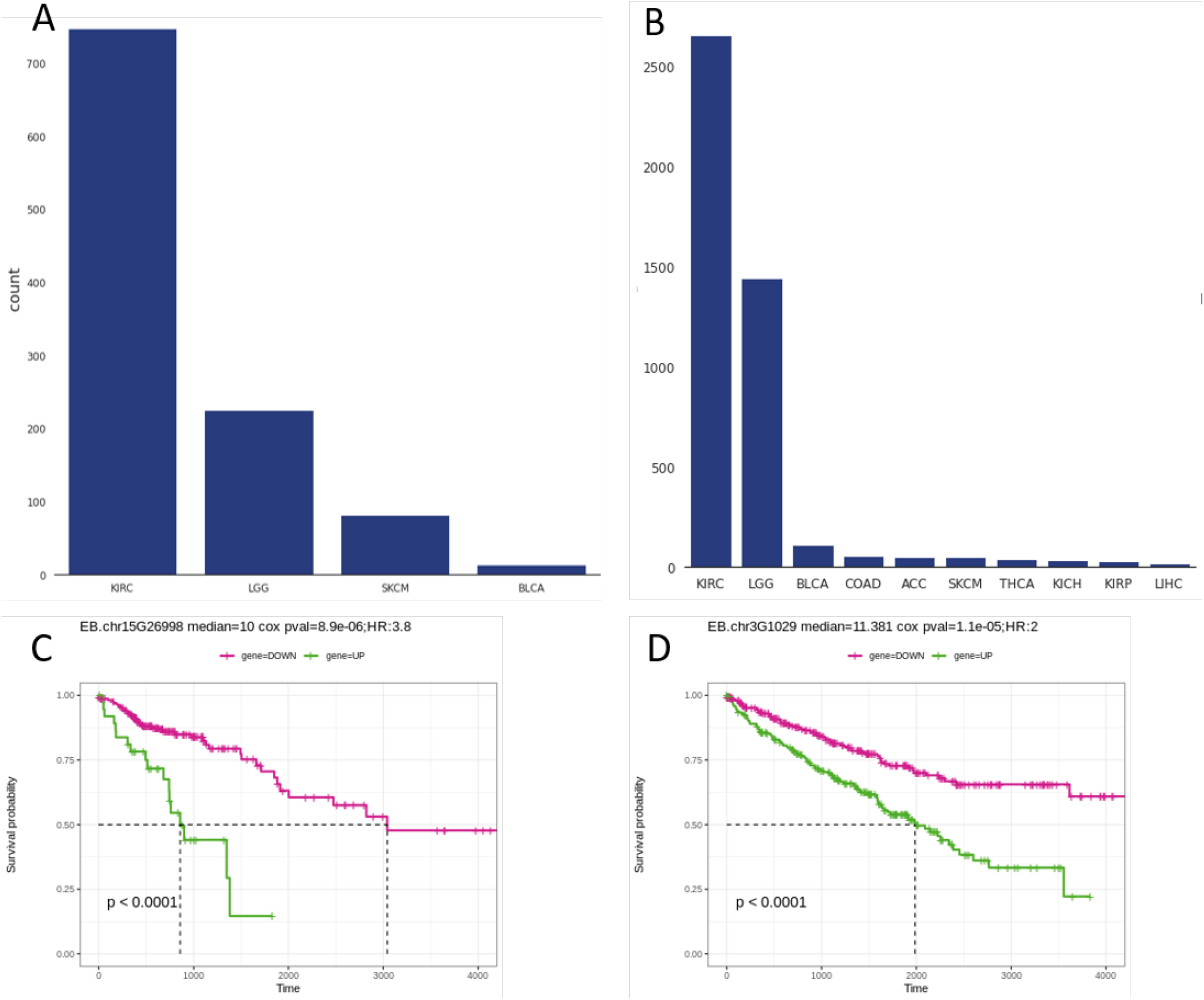
Multiple cancers express novel heEB genes that are associated with deleterious mutations and overall survival. Using Cox regression analysis, we labeled genes significantly associated with survival (adjusted p-value *<* 0.05). **A** heEB orphan genes that are upregulated in cancer and associated with unfavorable prognosis. **B** heEB orphan genes that are downregulated in cancer and associated with unfavorable prognosis. **C** Kaplan-Meier plot for orphan gene EB.chr15G26998. High expression of this gene is associated with poor survival in COAD. **D** Kaplan-Meier plot for orphan gene EB.chr3G1029. High expression of EB.chr3G1029 is associated with poor survival in KIRC.

### Novel heEB genes harbor millions of disease variants and are associated with overall survival

To investigate the potential involvement of EB genes in human diseases, we quantified the extent to which deleterious variants are represented in these previously unidentified transcripts (Supplementary Figure 9; Supplementary File 1).

**Fig. 9.**
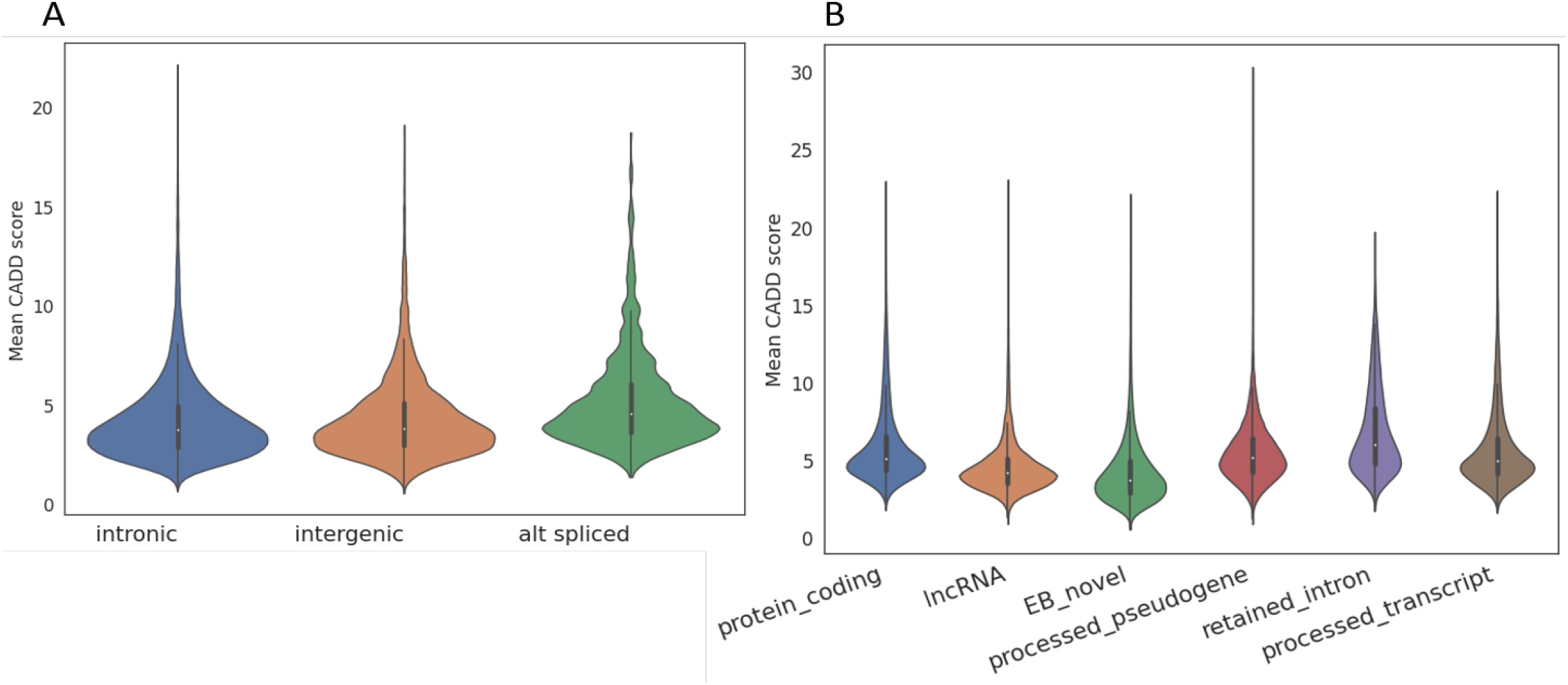
Distributions of Combined Annotation Dependent Depletion (CADD) scores across annotated genes and heEB genes. The CADD score corresponds to the deleteriousness of a variant in the human genome. Violin plots show the medians and distributions of the CADD scores. **A**. Intronic, intergenic and alternatively spliced EB genes. **B**. Annotated transcript types and heEB transcripts.

EB genes harbor a total of 327,565 COSMIC variants; 169,283 of these variants are predicted to be pathogenic. Skin, large intestine and lung contain particularly large numbers of pathogenic variants located within EB genes (Supplementary Figure 10).

**Fig. 10.**
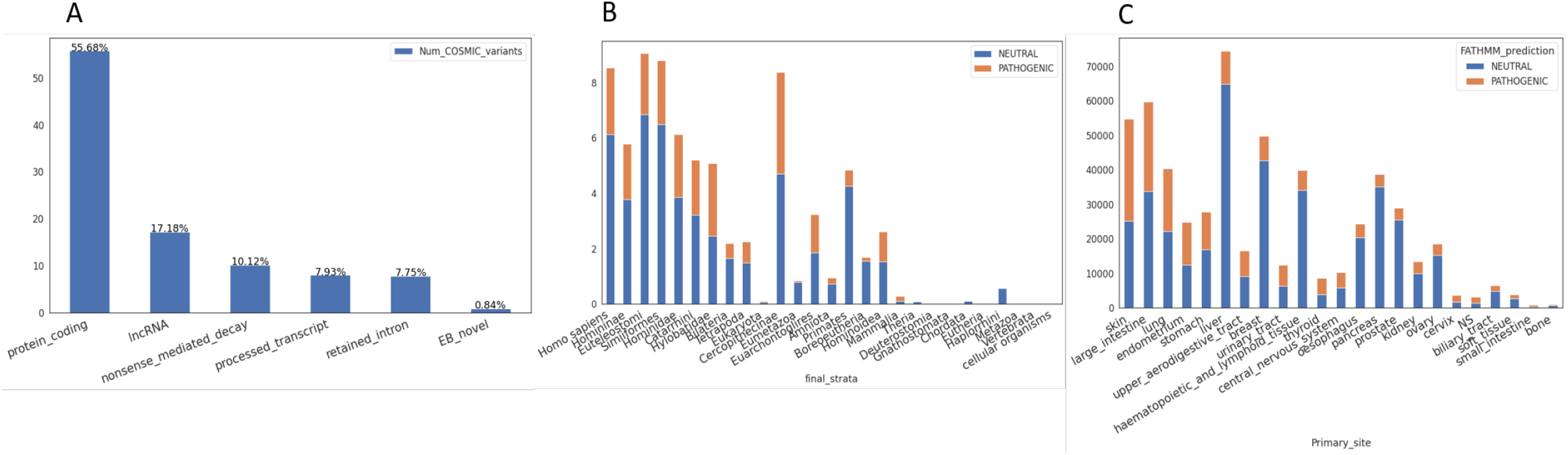
Distribution of COSMIC variants and FATHMM predictions for annotated trasncripts and for and high expression-evidence-based (heEB) transcripts. We queried the COSMIC non-coding variants that overlap with annotated and heEB transcripts. Number of COSMIC variants in: **A**. Classes of annotated and heEB transcripts; **B**. heEB genes grouped by phylostrata and colored by FATHMM predictions of pathogenic variants. Each bar represents number of pathogenic or neutral variants per transcript in the phylostrata. **C**. heEB genes grouped by tissue site and colored by FATHMM prediction for pathogenic variants.

Thousands of DE heEB genes show a significant association with overall cancer survival. The tumor types with the greatest number of survival-associated heEB genes are shown in Figure 8. The higher number of heEB orphan genes that are associated with favorable prognosis might partly be related to the higher number of heEB orphans that are downregulated in cancers.

### Evidence of peptides encoded by EB genes

To identify translated heEBs, we processed 289 Ribo-Seq samples from 23 independent studies and assessed the translation of all the annotated and heEB transcripts in these samples. A caveat in this analysis is that heEB genes tend to be sparsely expressed (5), and there was limited diversity in the Ribo-Seq samples relative to the 26,000 samples we processed to identify EB transcripts.

Using the ribotricer (43) tool, we identified 943 heEB transcripts with evidence of translation (Figure 21). Transcripts annotated in Gencode as “retained intron” (23,510 transcripts), “nonsense-mediated decay” (15,081 transcripts), “pseudogenes” (3,431 transcripts), and lncRNAs (13,272 transcripts) were also detected as translating novel ORFs (Figure 21).

We further investigated the phylostrata and transcript class of the heEB genes with translation evidence. The majority of heEB genes with evidence of translation (646) are human-specific orphan genes. However, two intergenic heEB genes with translation evidence are assigned to the oldest phylostratum, i.e., cellular organisms (Figure 21).

### Inferring functions of heEB genes

Co-expressed clusters of genes have been a successful approach to assign functions for genes about which little or nothing is known (44–47). However, because orphan genes are relatively young, many may not have fully developed regulatory mechanisms (2). To explore whether a guilt-by-association approach could provide a mechanism to develop hypotheses on possible functions of some heEB orphan genes, we analyzed 2,630 GTEx samples from brain. A Spearman correlation matrix was computed from the combat-seq (48) corrected counts data. We generated gene co-expression networks from the correlation matrix, selecting for highly correlated gene pairs above a threshold of 0.9 to construct a network that exclusively retained the strongest associations. We then partitioned the network into clusters using the MCL algorithm (49), resulting in 156 clusters of over ten genes; thus, each gene in a cluster had a correlation of 0.9 or greater to at least one other gene in that cluster. Some clusters contained predominantly orphan genes, other predominantly Gencode-identified protein-coding genes, and others a mixture of the two (Supplementary Data). We calculated Gene Ontology enrichment for each cluster with more than 10 transcripts with annotated functions.

A central driver of cellular regulation, particularly in the brain, is oxidative respiration (50), a process whose genes may be broadly co-expressed (47). We identified one co-expression cluster, Cluster 4945, that was highly enriched in oxidative respiration (”*proton motive force-driven ATP synthesis*”, adj pValue 1.4E-50) (Fig. 11). Cluster 4945 contains transcripts whose encoded proteins participate in the structure and function of the respiratory complexes, enable transport across mitochodrial membranes, and regulate the delicate balance between oxidative respiration and mitochondrial defense-response. Cluster 4945 contains two heEB transcripts: EB.chr3G17848 and the orphan gene EB.chr19G20849.

**Fig. 11.**
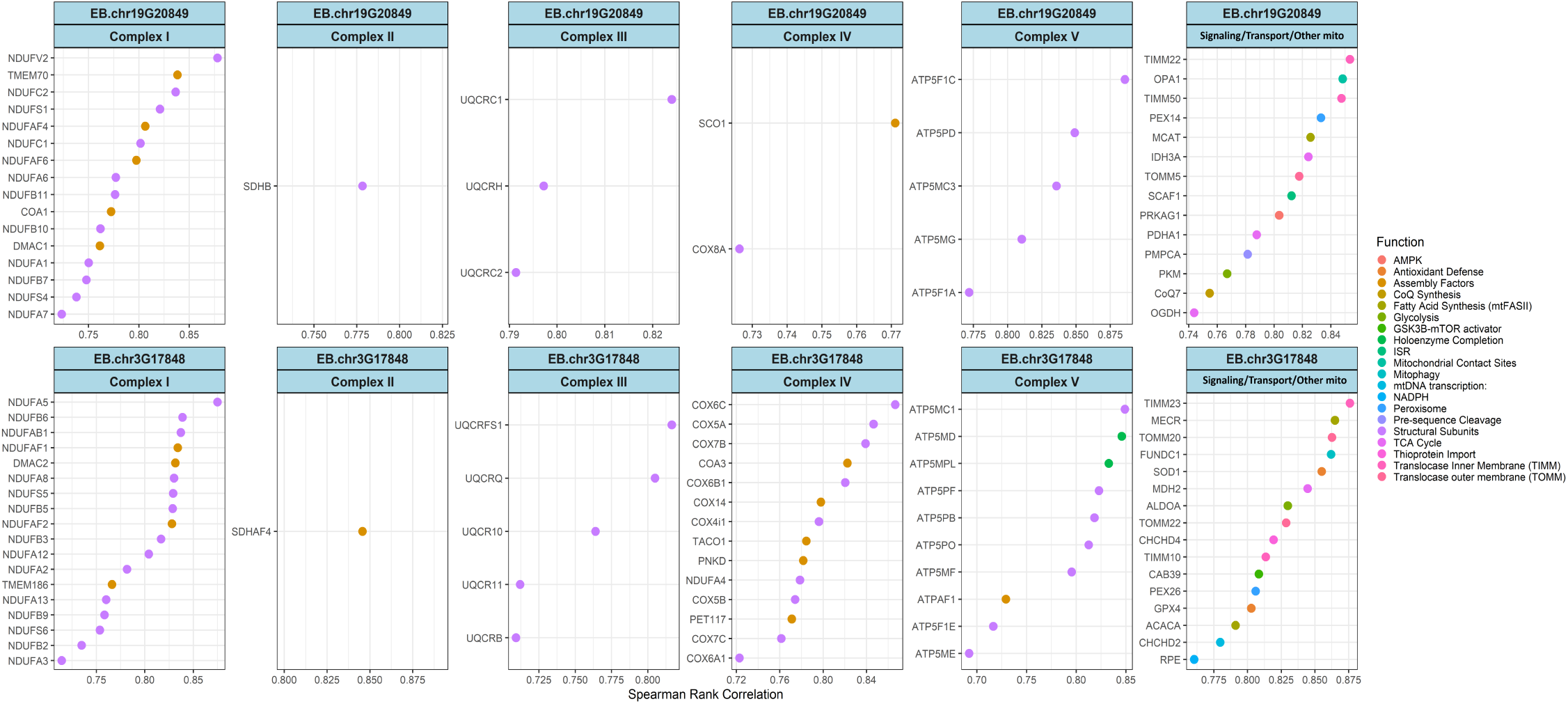
A cluster enriched in oxidative respiration contains two novel orphan genes. Cluster 4945 was obtained by MCL partitioning of a pairwise Spearman correlation matrix. The cluster contains the orphan gene EB.chr19G20849 the non-orphan heEB EB.chr3G17848.

**Fig. 12.**
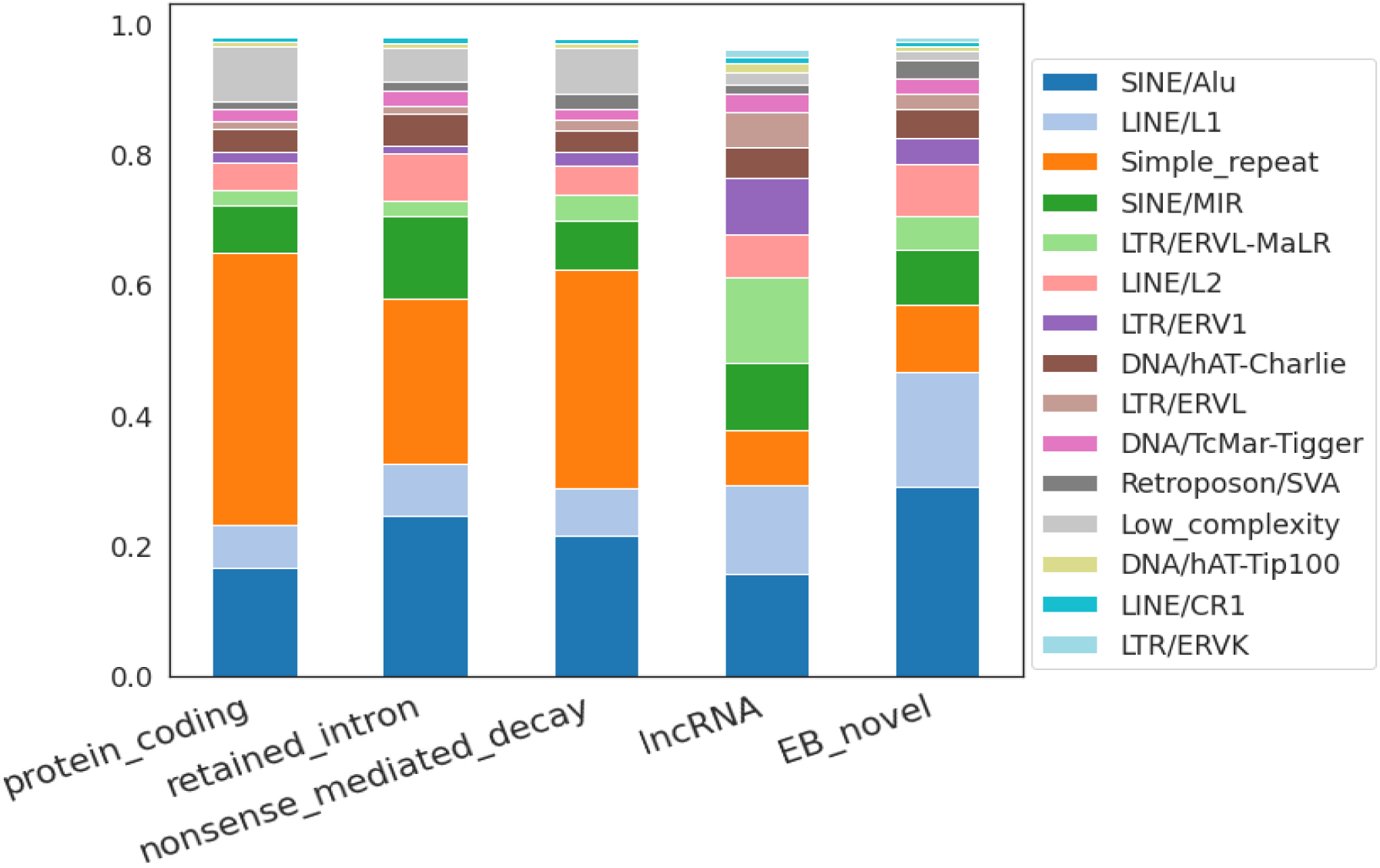
heEB transcripts have higher proportions of SINE/LINE repeats than annotated protein-coding transcripts. Repeatmasker was used to evaluate the repeats in annotated and heEB transcripts. SINE/Alu repeats are most prevalent class of heEB transcripts, versus “simple repeat” in protein-coding annotated genes.

## Discussion

A significant proportion of RNA molecules in a given biological sample is not annotated in community databases. We refer to this unannotated, yet expressed set of transcripts as the dark transcriptome (26). It includes coding and non-coding genes that have eluded annotation, sequence that is fodder for the evolution of new genes, and non-specific “noise” (26, 51– 55). Our premise is that within this dark transcriptome lies hundreds or thousands of as-yet-unannotated protein-coding orphan genes.

Although initially, almost all transcripts other than already-annotated genes were considered “noise” (51–55), recently, such transcripts are often annotated as ncRNA, despite that some of these “ncRNAs” have ORFs and identified functions for the encoded protein (11, 21, 24). Our study identifies novel transcripts that are *highly* expressed, with a special effort to retrieve those that are *selectively* expressed in diverse tissues, tumors, across sex and ancestries and may not be detected by other approaches (Figure 1; Supplementary File 3). Our data shows that over 80% percent of the total transcript accumulated in tissues is composed of unannotated transcripts. On the general principle that cells do not spend precious energy synthesizing vast amounts of useless material, we posit that many of these highly accumulated RNAs have functions, ranging from providing general scaffolding for nuclear function to having unique roles in biological processes.

Leveraging diverse tumor and non-diseased tissues, as well as COVID-19 infected tissues, we reprocessed data from over 28,000 RNA-Seq runs to capture novel high-expression transcripts associated with particular tissues, diseases, and ancestries. We supply the gene models for these heEBs as a UCSC Genome Browser track https://genome.ucsc.edu/s/jahaltom/Orphan%20Genes.

Because we partitioned our analysis into biological groups, we captured highly accumulated transcripts that would have been missed if we had combined median expression values across all conditions. For example, if we had pooled all GTEx samples *before* we filtered the RNA-Seq data for low expression, only 763 heEB transcripts would have been identified as highly expressed.

Several studies, including the largest previous to the current research (51), removed from consideration as novel genes those transcripts with single exons or no similarity to annotated genes in other species. Because a high proportion of orphan genes are monoexonic (2, 6) and, by definition, all are species-specific, most would have been eliminated. Notably, we provide a reproducible pipeline to identify orphan genes and other novel genes across as many as tens of thousands of samples and multiple conditions. The resultant catalog of the 79,023 highly and selectively expressed novel genes together with the annotated human transcripts can serve as a basis to decipher human diseases, novel cell types and complex gene evolutionary processes.

The genetic instability associated with cancer results in the creation of new ORFs, and is thought to account for transcripts expressed uniquely in cancers (56). Cancer- and patient-specific expressed neo-peptides are promising targets for cancer immunotherapies (57, 58). Interestingly, the aberrantly expressed transcripts from the “non-coding” genome offers a broader array of targetable neoantigens spanning different cancer types (57). While not explored in this study, certain tumor-specific heEBs may hold potential as targets for immunotherapy or for cancer vaccination. While other novel heEB genes associated with overall patient survival may provide promising candidates for diagnostic and therapeutic intervention.

We have processed over 250 terabytes of RNA-Seq data to detect millions of innominate transcripts in the molecular milieu of human tissue. By analyzing these data with respect to tissue, disease, ancestry and sex, we have identified over seventy nine thousand transcripts that are selectively-expressed at levels higher than the median annotated protein-coding gene. We quantify these transcripts, and calculate coexpression clusters, phylostrata, and genomic context. The resultant data provides an invaluable resource for hypothesis generation and testing.

## Materials and Methods

### RNA-Seq Datasets

RNA-Seq data for determination of EB genes were from the two most comprehensive human RNA-Seq repositories, the Genotype-Tissue Expression portal (GTEx) (31) and The Cancer Genome Atlas (TCGA; https://www.cancer.gov/tcga). All the non-diseased samples from GTEx are polyA+ RNA-seq libraries. The library selection protocol differs for TCGA cohorts. Corresponding sample metadata were also obtained from GTEx and TCGA portals.

Stranded human RNA-Seq data were downloaded from NCBI-GEO accessions GSE69241 (35) and GSE146009 (40). Stranded chimpanzee RNA-Seq data was downloaded from GSE69241 (35). Single cell RNA-Seq data for human breast, liver, and testis tissues were obtained from accessions GSE164898 (59), SRP285767 (60) and SRP214255 (61) respectively. 289 samples of human Ribo-Seq data from 23 studies were obtained from NCBI-GEO and RPFdb (62) (Supplementary File 5). RNA-Seq samples related to COVID-19 were obtained from SRA https://github.com/jahaltom/Orphan-Gene-Supp.

### Evidence-based orphan gene annotation pipeline

We engineered a scaleable best-practice pipeline for detection of annotated and unannotated genes using 26,985 RNA-Seq samples from GTEx and TGCA. The abstract pipeline flowchart is shown in Figure 13; the steps are discussed in the following sections.

**Fig. 13.**
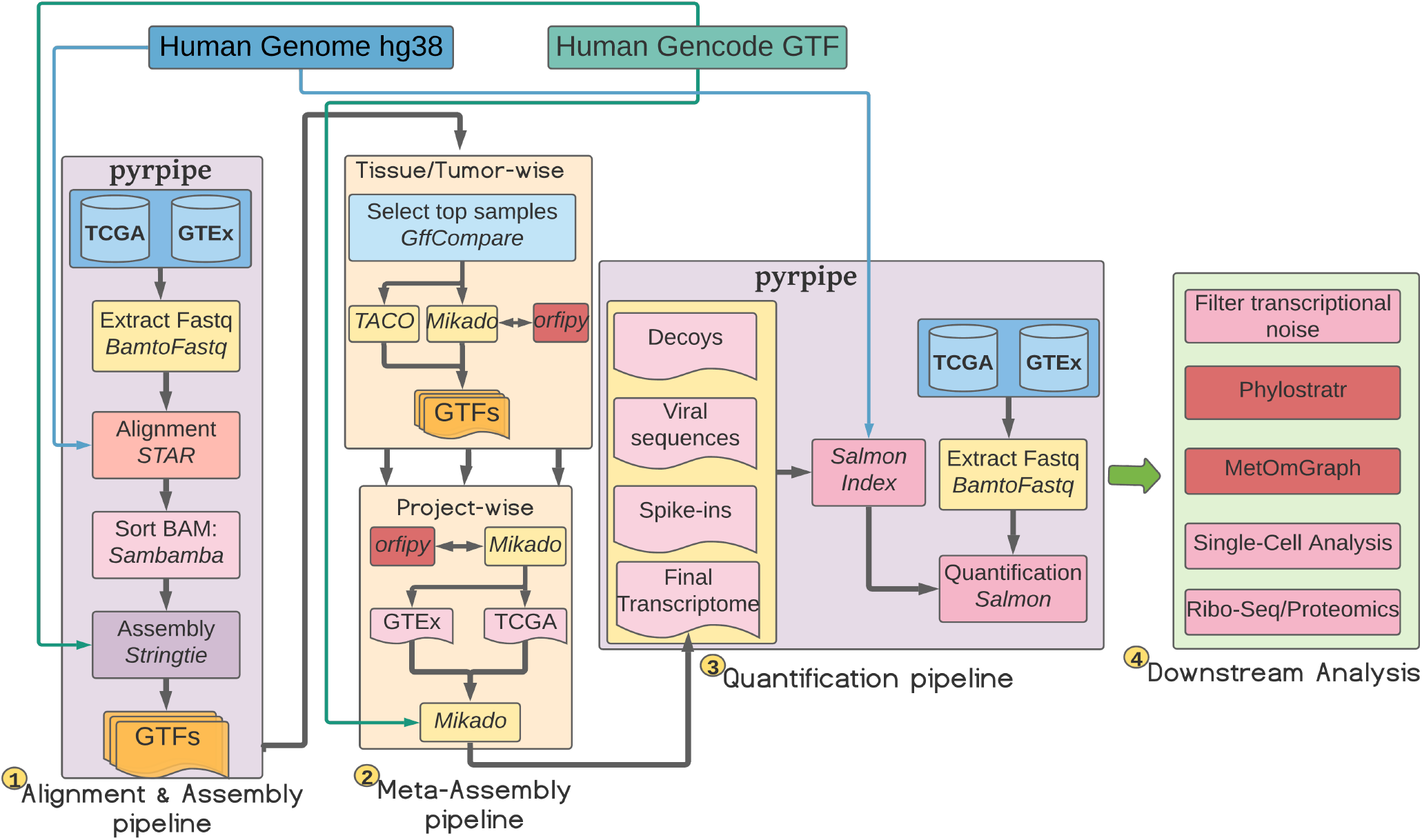
Workflow of the study. The alignment and quantification pipelines are implemented in pyrpipe (63). For alignment, BAM files from 26,985 from RNA-Seq samples from GTEx and TGCA] were converted to fastq using biobambam2. STAR (64) was run in 2-pass alignment mode to align reads to the human reference genome (GRCh38 GCA_000001405.15). Transcripts were assembled using Stringtie (65). Individual transcriptomes were consolidated into single transcriptome using our meta-assembly pipeline, consisting of Mikado (66) and Taco (67). ORFs were identified by orfipy (68). For quantification, a Salmon index was build using human annotated and novel EB transcripts, with whole human genome sequence used as a decoy. GTEx and TCGA fastqs were passed to Salmon’s *quant* function for quantification. phylostratr (36) was used to infer phylostrata of all Gencode-annotated proteins and for each EB transcript.

#### Read alignment and transcriptome assembly

The RNA-Seq raw reads from GTEx and TCGA samples were extracted from the downloaded BAM files using biobambam2 (69). Only samples with paired-end layout were considered. The RNA-Seq reads were aligned to the human reference genome (version GRCH38 GCA_000001405.15) using STAR (64). Transcripts were assembled using Stringtie (65) with reference GTF annotation as *guide*. We implemented the full RNA-Seq alignment and assembly processing pipeline using pyrpipe (63) and ran it on PSC Bridges HPC. The GTEx processing pipeline is available from github.com/urmi-21/pyrpipe/blob/master/case_studies/GTEx_processing.

#### Meta-assembly pipeline

From the assembled transcriptomes for each tissue/tumor/disease type, we considered a maximum of the 200 samples that were richest in expressing unannotated EB genes. We used gffcompare (70) to compare the assembled transcripts with the reference annotation and select samples that have highest unannotated transcripts.

A consolidated reference annotation was built using a combination of Taco (67) and Mikado (66) tools. Taco was run to retain transcripts with a minimum 100 nt in length with expression based filter turned off. Mikado was tuned to identify transcripts longer than 100 nt. A custom Mikado *scoring* file was used to enhance the prediction of novel transcripts. We used orfipy (68) to find all possible ORFs of 99 nt or longer in the transcripts. orfipy results were provided to mikado *serialise* step to annotate the CDS and other features.

First, Mikado and Taco were run independently to merge annotations from the same tissue and tumor type. Then, individual annotations from all non-tumor tissues were merged and the same was done for all tumors. We used gffcompare to compare tumor and non-tumor Taco annotations with corresponding Mikado annotations. Novel transcripts not predicted by Mikado were added to get final GTEx and TCGA annotations. Finally, we used mikado to merge the GTEx, TCGA, and Gencode reference annotations to arrive at a single consolidated GTF file (Figure 13). This annotation file contained 1,384,542 transcripts.

#### Pre-screening of consolidated assembly

We used Mikado’s compare utility to compare our annotation with the Gencode reference annotation. We first removed all the transcripts from our annotation that matched the reference annotation leaving only novel transcripts. We removed fusion products, and transcripts overlapping with annotated transcripts except for those marked as *alt-spliced*. Thus retained set contained novel alt-spliced, intronic, and intergenic transcripts.

We selected a subset of alt-spliced transcripts by running phylostratr (choosing a smaller set of reference species for quick analysis) (36) to retain only those that were human-specific. Then, we ran cd-hit (71) on the transcripts (90% similarity over 90% length) to remove redundant transcripts.

For quantifying gene expression across samples, we added the alt-spliced, intronic, and intergenic transcripts to all Gencode-identified transcripts.

#### Transcript expression quantification pipeline

We used Salmon’s selective alignment approach (72, 73) to quantify all the novel EB and annotated transcripts. Human reference genome along with viral sequences were used as decoys. Final transcript level counts were aggregated to gene level counts using custom python script.

#### Phylostratigraphy and genomic context

We used *phylostratr* (36) to infer phylostrata of ORFs of all Gencode annotated transcripts and all EB transcripts that contained an ORF of length >99 nt (33 aa). The annotated proteomes for 241 species binned into 26 phylostrata were used for phylostratal inference (Supplementary File 4). *phylostratr* automatically selects species for a balanced representation of phylostrata, retrieves protein sequences from these species, infers phylostrata and returns diagnostics.

Because *phylostratr* relies on known protein-coding genes (default-mode), some human genes might be predicted as orphans simply because they are not yet annotated in genomes of related species. To account for as-yet unannotated genes in comparison species, we used Liftoff (37) to map all heEB transcripts to five ape species: chimpanzee, gorilla, macaque, gibbon, orangutan; and four other species: cattle, zebrafish, mouse and rat. Liftoff mapped 13,921 EB transcripts to genomes with ORFs that encoded for proteins with similarity. The final phylostratal assignments of EB genes were based on *phylostratr* as refined by Liftoff.

### Filtering novel genes based on level of expression

We retained a total of 54,794 novel EB transcripts (54,748 genes) from the GTEx and TCGA datasets, selecting for expression above the median of expression of annotated protein-coding genes, according to the Filtering Algorithm below. (A less stringent criteria to filter would be to use annotated long non-coding genes instead of annotated protein-coding genes in Step 4).

Filtering Algorithm:

1. Group samples by tissue sampling site, race, and sex
2. Discard groups with less than 5 samples
3. Within each group, compute median of all transcripts and reduce the data to keep all the transcripts with median >= 1 TPM.
4. Compute the median of the median of all remaining annotated protein-coding transcripts
5. Filter out EB transcripts with median expression less than the median computed in step 4

Code used in the analysis is well-documented in GitHub (https://github.com/urmi-21/Human_orphan_genes).

### Intronic transcripts coexpression analysis

To investigate the coexpression patterns of intronic transcripts with the corresponding annotated transcript that contains the intron, we computed pairwise Spearman correlations for each intronic-annotated transcript pair. The correlation values were computed using the raw counts in a tissue/tumorspecific manner.

### Differential expression analysis

DESeq2 was used for differential gene expression analysis between TCGA tumor and GTEx undiseased samples (74, 75). We controlled for age, sex and race. For gender-specific tissues, only age and race factors were considered. Samples with missing data on age, sex or race were removed from the analysis.

We identified DE genes between conditions using contrasts (http://bioconductor.org/packages/devel/bioc/vignettes/DESeq2/inst/doc/DESeq2. html#contrasts)according to DESeq2 model (74, 75) Differential expression criteria

> *>* 2*foldchange, adjustedp − value*

< 0.05.

### Survival analysis

For each tumor type, DE genes were analyzed in relation to survival of the individuals. The data for overall survival was obtained from TCGA meta-data. Specifically, the columns *days_to_death* and *days_to_last_follow_up* are used.

We used the R package, survival (https://cran.r-project.org/web/packages/survival/index.html), to access the survfit and coxph functions for survival analysis. We used survfit to plot Kaplan-Meier survival curves, and coxph to fit a Cox proportional hazards regression model, while controlling for the effect of age, sex, and race.

P values for the Cox regression model were adjusted using the Benjamini-Hochberg method. Genes with adjusted p-value *<* 0.05 are reported as significantly associated with overall survival.

### Processing stranded RNA-Seq

Human stranded RNA-Seq data were obtained from two studies: GSE146009 and GSE69241. The stranded RNA-seq data were run through out salmon quantification pipeline to get the estimated read counts for annotated and EB genes.

Chimpanzee reference genome and transcriptome were obtained from GCA_002880755.3; chimpanzee stranded RNA-Seq data was downloaded from accession GSE69241. To the reference transcriptome, we added 11,510 human EB transcripts that were annotated as being in the chimpanzee genome by Liftoff. We used salmon to quantify the annotated and EB transcripts. The quantification pipeline was implemented using pyrpipe (63).

### Single-cell RNA-Seq data processing

Single-cell datasets were downloaded from accessions: SRP301923 (breast), SRP285767 (liver), SRP214255 (testis), All single-cell data used were sequenced using the 10X Genomics’ Chromium™ platform. We processed processed these samples using the 10X Genomics’ Cell Ranger version 6.0.1 (76). We created a custom reference transcriptome to be used with Cell Ranger by running *mkref* command with all the annotated and highly expressed EB genes. Cell Ranger’s *count* command was run each RNA-Seq sample with custom reference created with *mkref*.

Downstream analysis used scanpy (77, 78) in a study-specific manner. All samples from a study were concatenated into a single *AnnData* object. Data pre-processing steps included removing cells with too many mitochondrial genes expressed or too many total counts and log normalizing the data.

Genes used for single cell identification in lung were from (79–82). Visualization of lung data was using Morpheus, https://software.broadinstitute.org/morpheus

### Ribo-Seq analysis pipeline

We used Ribosome profiling (Ribo-Seq) data from 23 studies and processed a total of 289 samples. Ribo-Seq data were processed on a sample-by-sample basis. Sequencing adapters were removed before aligning the reads to the human reference genome using STAR aligner (64). We used ribotricer (43), run with suggested parameters for human samples., to detect translating ORFs in all the annotated and highly-expressed EB transcripts.

### Variant analysis

We queried gnomad ((v2.1.1 liftover data) (83) to identify variants in the annotated and EB genes, identifying 6,985,430 variants located within the 54,794 heEB transcripts. Then, we searched disease causing mutations as described in the Catalogue Of Somatic Mutations In Cancer (COSMIC) (84), specifically, COSMIC’s “non-coding variants”, defined as variants that occur within intronic or intergenic regions of the genome (https://cancer.sanger.ac.uk/cosmic/help/ncv/overview). Tabix (85) was used to query the variants from the zipped vcf files. Deleteriousness was examined using Combined Annotation Dependent Depletion (CADD) scores (86). Mean CADD score for each transcript was calculated using custom python and bash scripts (Fig. 9).

### Computation of co-expression clusters and annotation of genes

Gene co-expression networks were computed from a Spearman’s correlation matrix, in a tissuetumor-wise manner, on ComBat-seq corrected raw counts (48) at a threshold of 0.9. MCL algorithm (49) was used to identify gene-clusters in the network. The best partition of the network was selected based on the maximum modularity value. Functional enrichment of clusters was determined by ToppGene (87).

### Repeat masking analysis, computation of protein disorder and other features

RepeatMasker (http://www.repeatmasker.org) was used to identify repeats and low-complexity regions in all the annotated and EB transcripts. RepeatMasker was run with the *species* parameter set to human.

Protein disorder was predicted using IUPred2A (88) in both short and long modes. Emboss (89) was used to calculate the isoelectric points. Custom python script was written to compute CDS length, GC%, Codon Usage (CU), and Relative Synonymous Codon Usage (RSCU).

## Supplementary Data

Supplementary data and files are available from https://github.com/urmi-21/Human_orphan_genes. Genomic context of highly expressed EB genes is at UCSC genome browser https://genome.ucsc.edu/s/jahaltom/Orphan%20Genes. Project files to inter-actively explore the COVID-19 data using MetaOmGraph are available at https://iastate.box.com/s/u3a9fuwqo7h3cpfzs9b9w1b5g5jxqv3x.

## ACKNOWLEDGEMENTS

We are grateful to all members of the Wurtele lab and to our COV-IRT (https://www.cov-irt.org/) colleagues, particularly Stephen B. Baylin and Michael J. Topper, for helpful and stimulating discussions. We thank Luca Venturini and other developers of Mikado, for providing support for their tool. We thank Robin Gogerty, Jake Miller, and Levi Baber at Iowa State University for help with administrative and IT services. This work is funded in part by the National Science Foundation award IOS 1546858, “Orphan Genes: An Untapped Genetic Reservoir of Novel Traits” and by the Center for Metabolic Biology, Iowa State University. This work used the Extreme Science and Engineering Discovery Environment (XSEDE), which is supported by National Science Foundation grant number ACI-1548562. In particular, it used the Bridges HPC environment through allocations TG-MCB190098 and TG-MCB200123 awarded from XSEDE and the HPC Consortium.

## Supplementary Note 1: Supplementary Figures

**Fig. 14.**
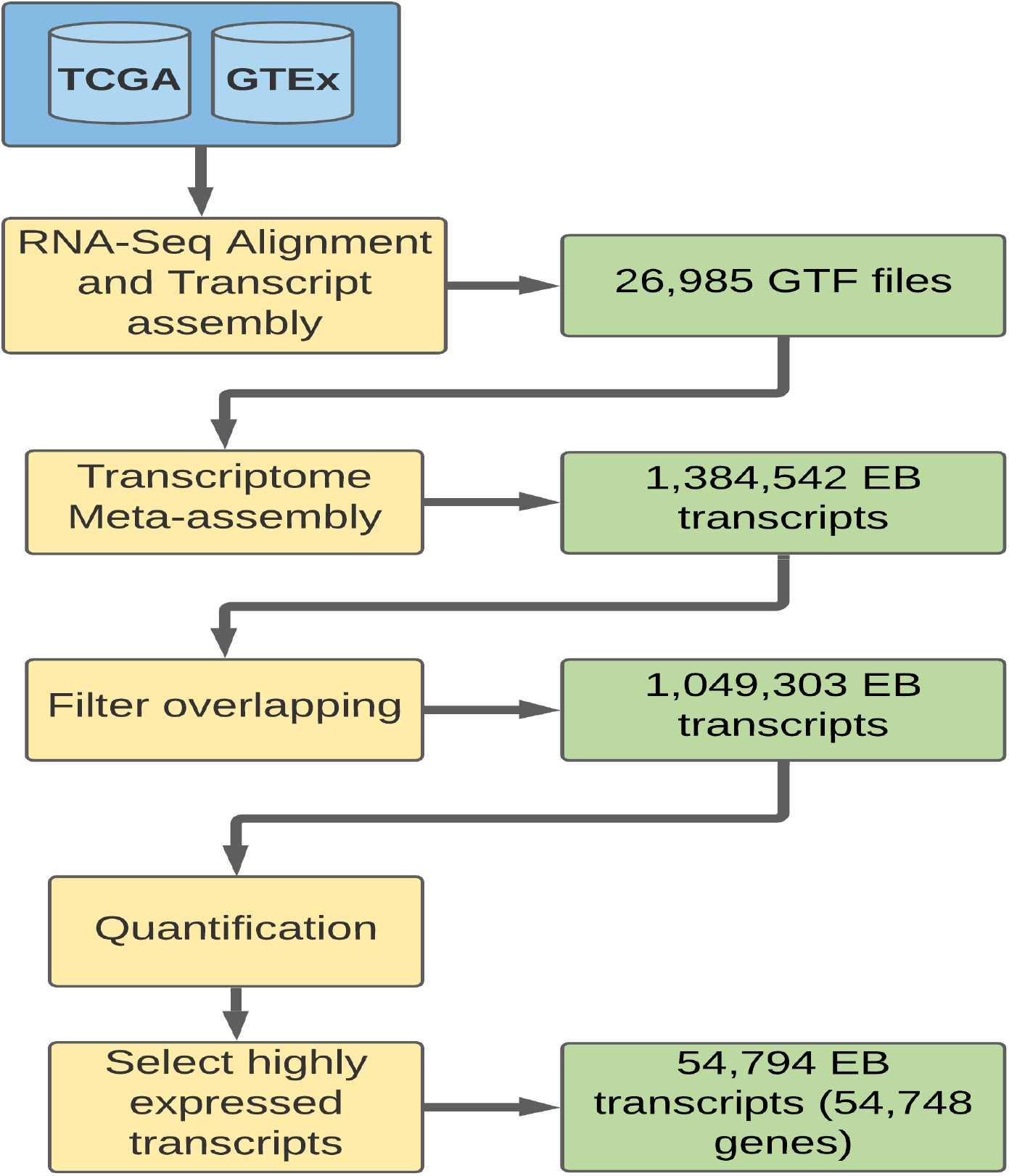
Number of EB transcripts identified at each step of the workflow used in this study.

**Fig. 15.**
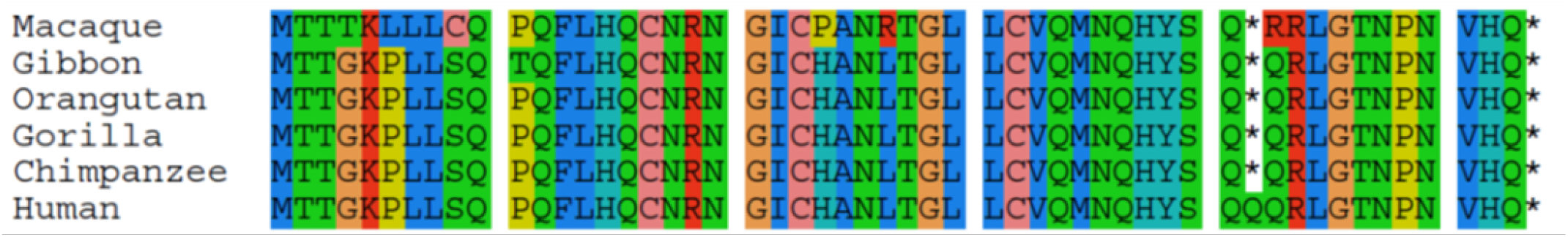
Example of an intergenic EB gene identified in this study. Liftoff mapped the gene EB.chr6G21600 to genomes of closely related species, in which the protein is terminated by an in-frame stop codon. This stop codon is mutated in humans, thus the ORF codes for a longer protein (53 AA).

**Fig. 16.**
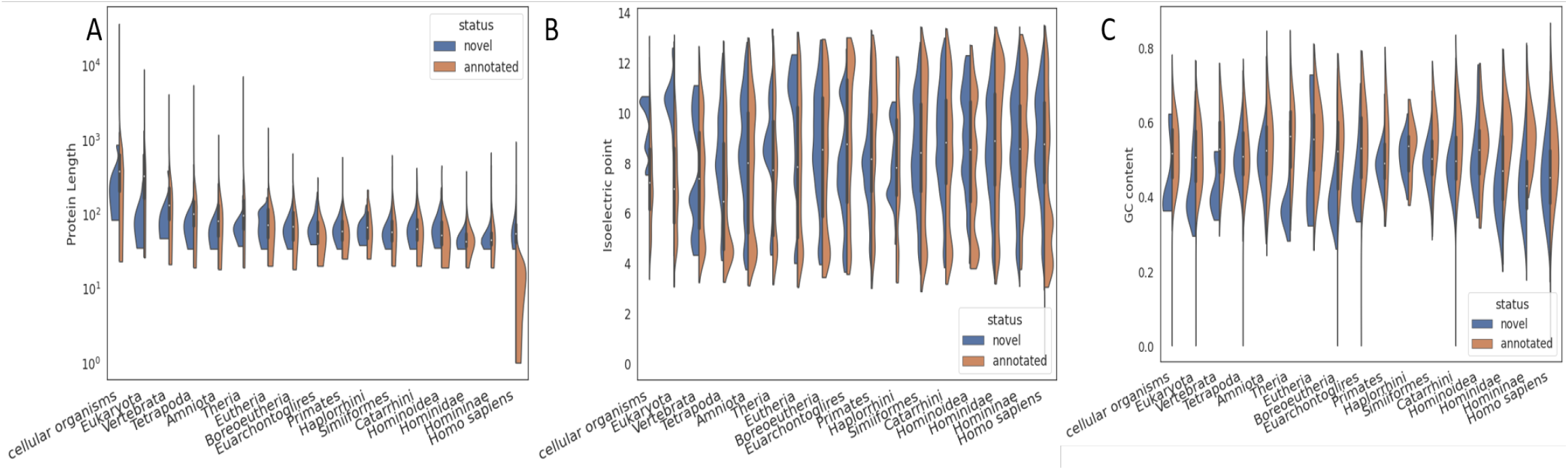
Comparison of features across phylostrata for proteins of annotated and EB genes. **A**. Protein length, **B**. Isoelectric point, **C**. GC content.

**Fig. 17.**
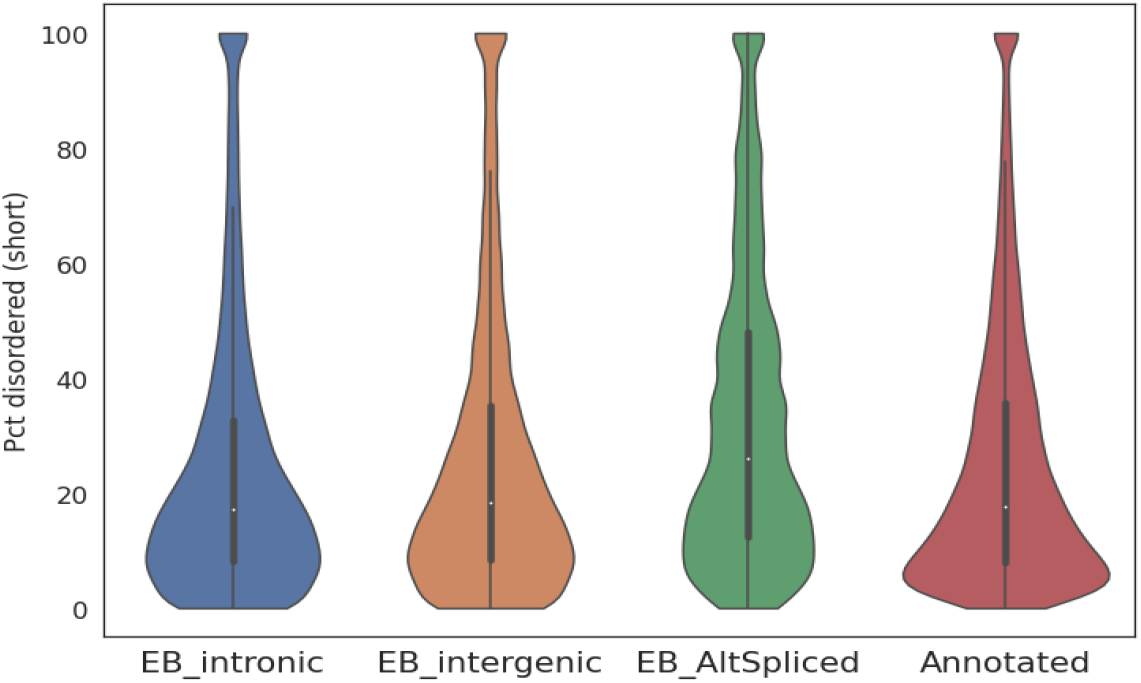
Distribution of percent disordered residues predicted for the heEB and annotated orphan proteins are similar to that of more ancient genes. Disordered residues within each protein were predicted using IUPred2A’s short disorder prediction (88). Proteins with high intrinsic structural disorder are unable to fold into aggregates and less likely to be harmful (90, 91). Several studies predict proteins of orphan genes to be highly disordered (2, 4, 90, 92, 93). However, the contrary has also been reported (94). We do not detect a significant difference in predicted disorder among proteins encoded by either the human novel heEB orphan genes or the Gencode-annotated protein-coding genes. Spliced heEB genes that overlap with annotated genes show a slightly higher median value for disorder, consistent with a previous study that reports young ORFs mapping to exons have elevated disorder (90). We surmise that these discrepancies in predictions of disorder among orphan genes can be attributed both to differences in the approach used to identify the orphans, and to the species analyzed.

**Fig. 18.**
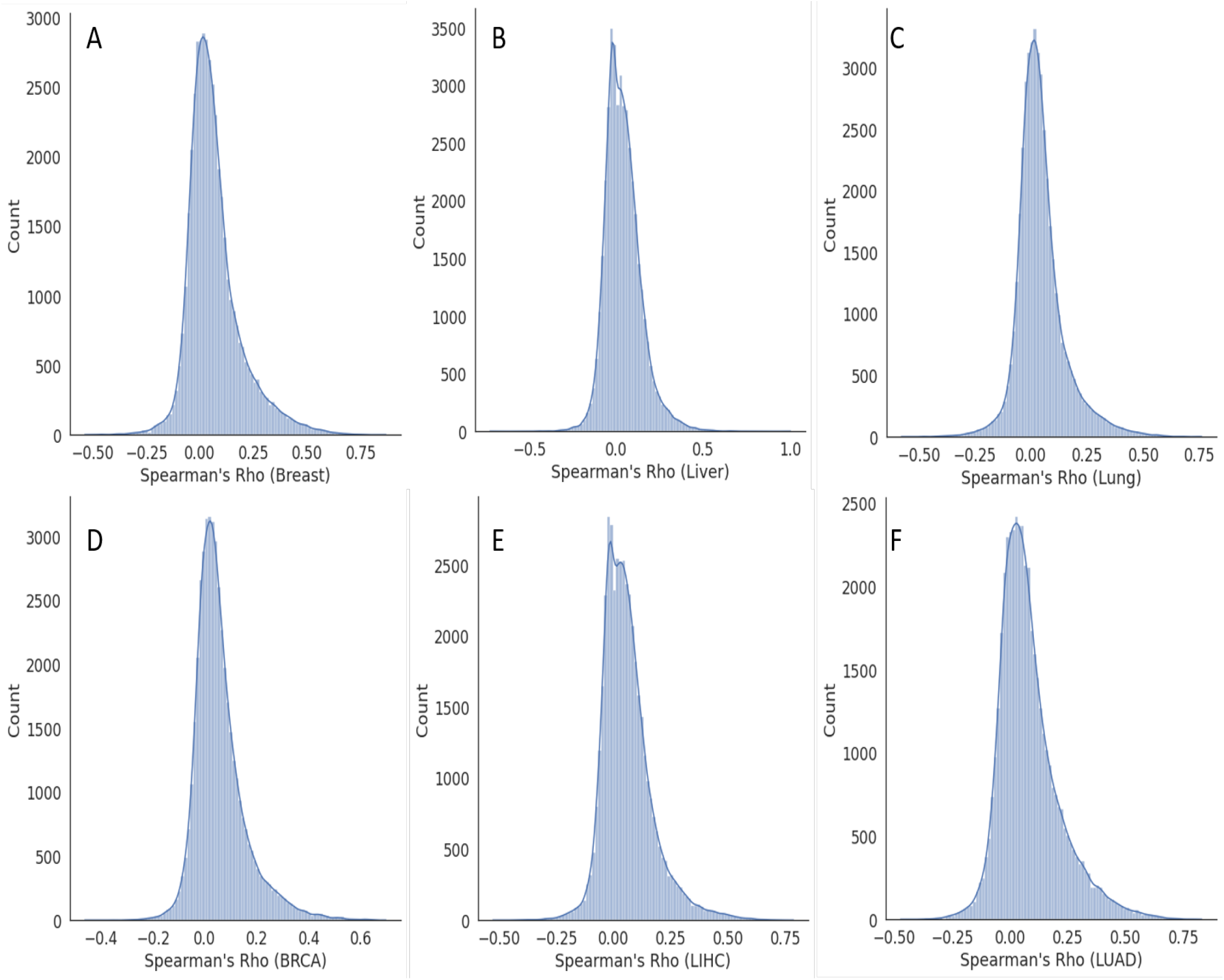
Distributions of Spearman’s correlation values between each intronic EB transcript and a randomly chosen annotated transcript. Distributions of Spearman’s correlation values for transcript pairs in: **A**. GTEx breast samples. **B**. GTEx liver samples. **C**. GTEx lung samples. **D**. TCGA BRCA samples. **E**. TCGA LIHC samples. **F**. TCGA LUAD samples. Some EB transcripts might represent processing errors, e.g., premature termination, intron retention, or intronic reads, though careful studies indicate many novel transcripts are intronic (34, 35). Reasoning that high expression correlation of an intronic EB transcript with its corresponding annotated transcript might reflect a processing artifact, we computed Spearman correlations for each intronic heEB transcript with the transcript that encompasses it, in a tissue specific manner. The mean correlation value was less than 0.3 (Figure **??**; Supplementary File 1). For comparison, mean correlation between an heEB and a gene chosen at random is 0.2 (Supplementary Figure 3 and File 1). These results indicate expression of intronic EB gene-annotated gene pairs are largely independent. Spearman’s correlations are computed for each pair, i.e, intronic EB transcript and corresponding randomly chosen annotated transcript, over samples for a tissue or tumor.

**Fig. 19.**
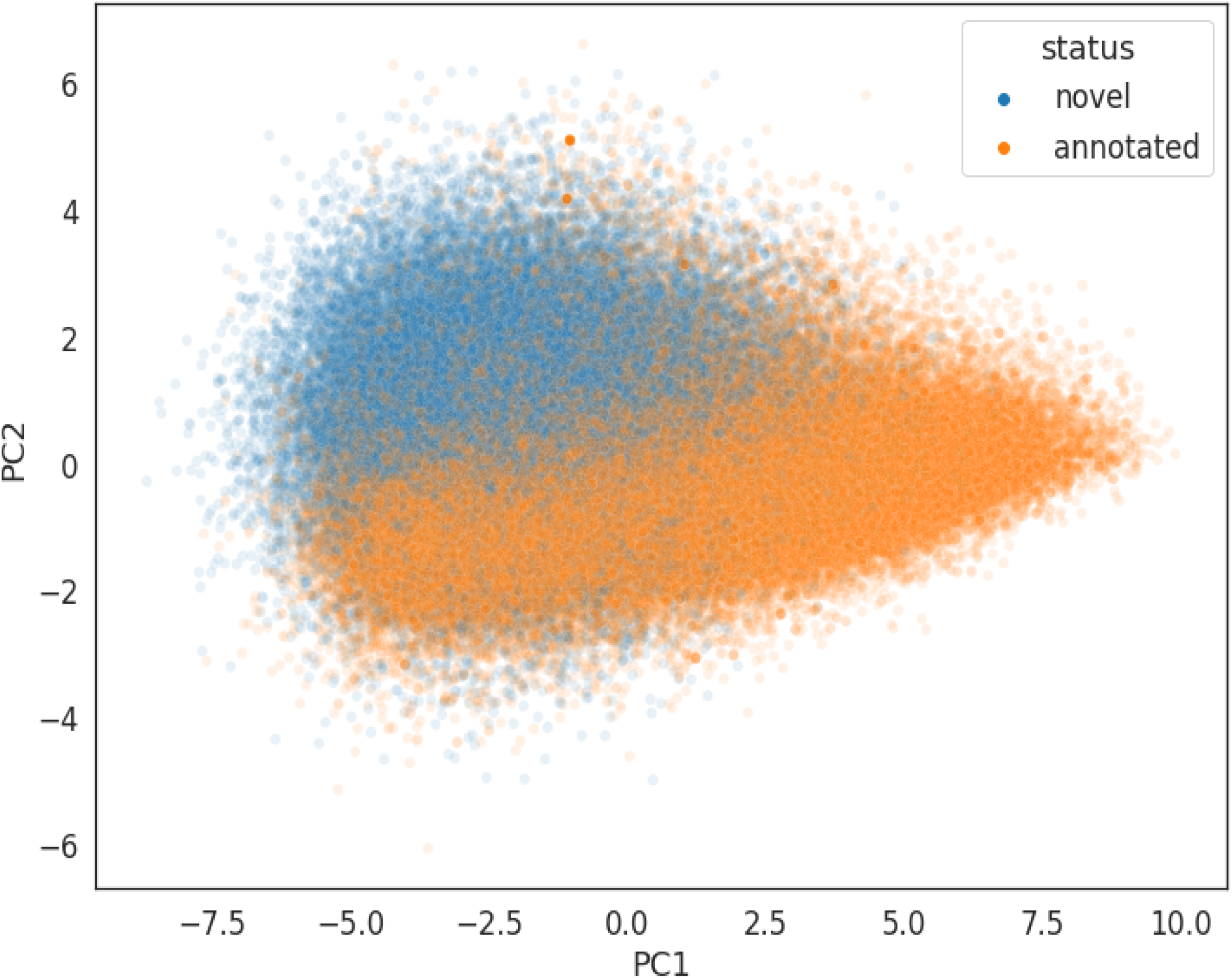
PCA plot using Relative Synonymous Codon Usage (RSCU) values of all pcEBs and annotated protein coding transcripts.

**Fig. 20.**
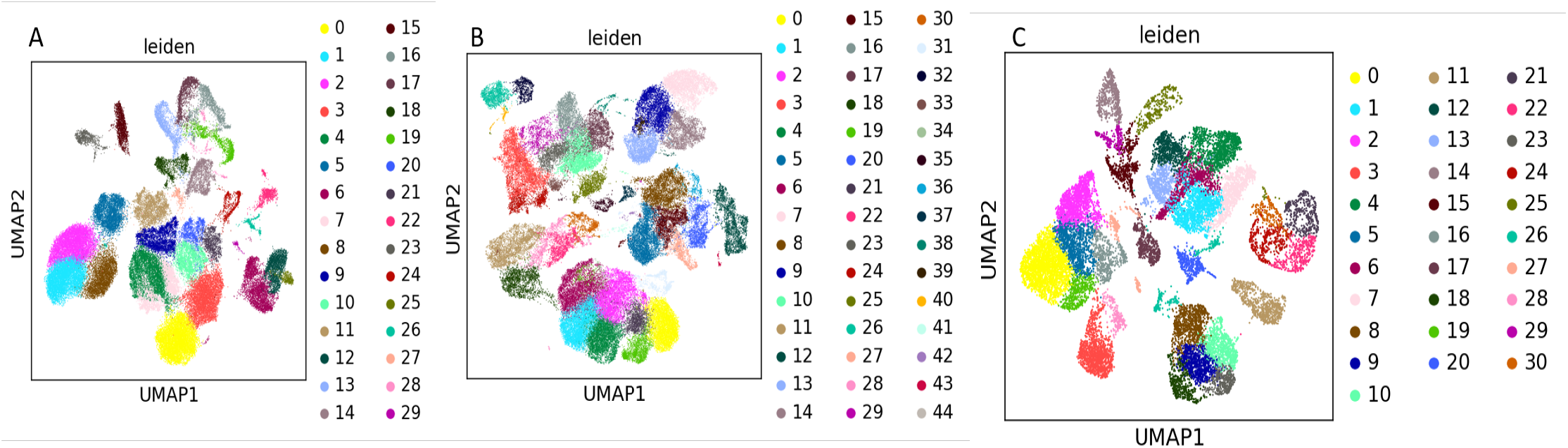
Single-cell RNA-Seq clusters identified in **A**. Liver **B**. Breast and **C**. Testis datasets.

**Fig. 21.**
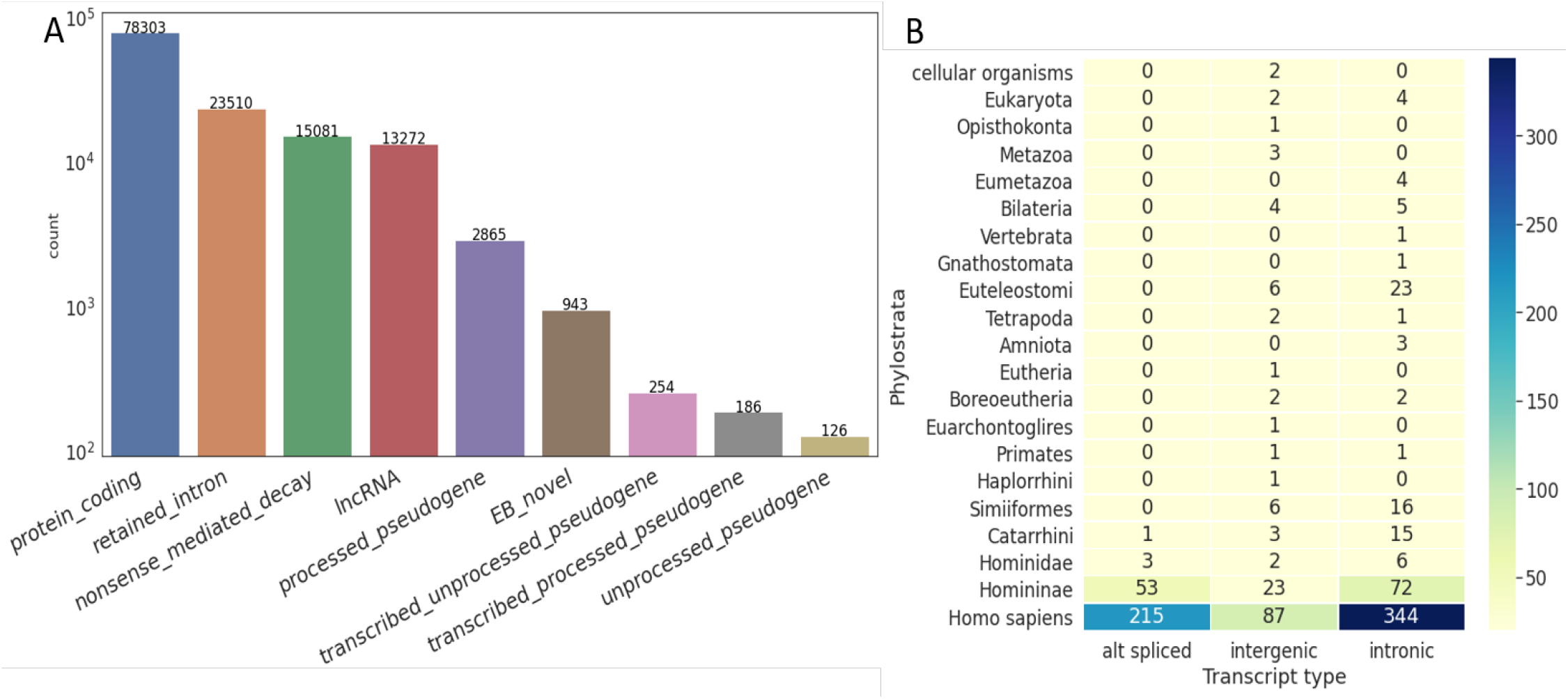
Translating ORFs in Ribo-Seq datasets. 289 Ribo-Seq samples in 23 studies were processed and translation of annotated and heEB transcripts assessed. **A**. Number of unique translating ORFs detected across various transcript types. **B**. Number of heEB transcripts with evidence in Ribo-Seq data, by phylostrata and transcript class.

**Fig. 22.**
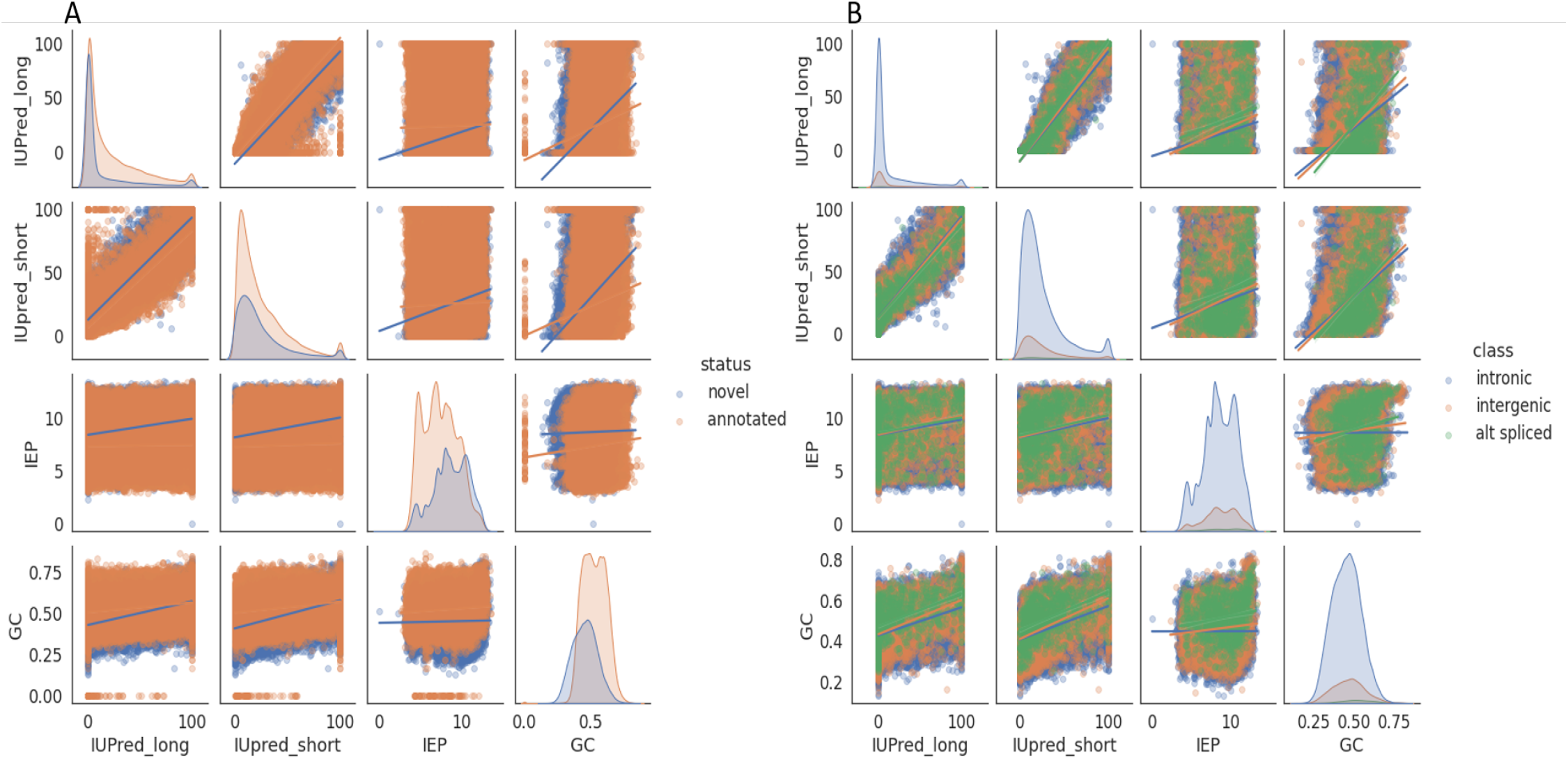
Variation of protein disorder (IUpred) and isoelectric point (IEP) with GC content. Lines are estimated by fitting a linear regression model. **A**. All EB and annotated proteins. Orange, annotated; blue, EB **B**. Only EB proteins. Orange, intergenic EB; blue, intronic EB; green alt-spliced EB.

## Notes

https://github.com/urmi-21/Human_orphan_genes/tree/main

### Competing Interest Statement

The authors have declared no competing interest.

### Summary of Updates

The revised manuscript corrects Author Affiliations

